# Effect of Sequence Depth and Length in Long-read Assembly of the Maize Inbred NC358

**DOI:** 10.1101/858365

**Authors:** Shujun Ou, Jianing Liu, Kapeel M. Chougule, Arkarachai Fungtammasan, Arun Seetharam, Joshua Stein, Victor Llaca, Nancy Manchanda, Amanda M. Gilbert, Xuehong Wei, Chen-Shan Chin, David E. Hufnagel, Sarah Pedersen, Samantha Snodgrass, Kevin Fengler, Margaret Woodhouse, Brian P. Walenz, Sergey Koren, Adam M. Phillippy, Brett Hannigan, R. Kelly Dawe, Candice N. Hirsch, Matthew B. Hufford, Doreen Ware

## Abstract

Recent improvements in the quality and yield of long-read data and scaffolding technology have made it possible to rapidly generate reference-quality assemblies for complex genomes. Still, generating these assemblies is costly, and an assessment of critical sequence depth and read length to obtain high-quality assemblies is important for allocating limited resources. To this end, we have generated eight independent assemblies for the complex genome of the maize inbred line NC358 using PacBio datasets ranging from 20-75x genomic depth and N50 read lengths of 11-21 kb. Assemblies with 30x or less depth and N50 read length of 11 kb were highly fragmented, with even the low-copy genic fraction of the genome showing degradation at 20x depth. Distinct sequence-quality thresholds were observed for complete assembly of genes, transposable elements, and highly repetitive genomic features such as telomeres, heterochromatic knobs and centromeres. This study provides a useful resource allocation reference to the community as long-read technologies continue to mature.

## Main

During the two decades following the publication of the first larger eukaryotic genomes (i.e., *Drosophila melanogaster*^1^ and *Homo sapiens*^2^), considerable progress has been made in sequencing technology and assembly methods, improving our basic knowledge of genome complexity across the tree of life. We now understand that genome composition (*e.g.*, gene complement, the extent of intergenic space, and the landscape of transposable elements (TEs)) varies substantially at both the inter- and intraspecific levels. For example, comparing the *Arabidopsis thaliana*^3,4^ and bread wheat (*Triticum aestivum*)^5^ genomes demonstrates a >100-fold difference in genome size (0.12 Gb and 14.5 Gb, respectively) and substantial variation in both gene number (32,041 versus 107,891 annotated gene models) and repeat content (21% versus 85%).

The goal of robust genome assembly is to capture and accurately represent all components of a genome so their biology may be accurately studied. Next-generation assemblies initially relied on short-read data due to cost and technological limitations. While these assemblies represented genes reasonably well, repetitive regions containing transposable elements and tandem repeats were either omitted or highly fragmented^6^. Newly developed long-read sequencing technology now enables contiguous assembly of even the repetitive fraction of eukaryotic genomes^7^ with, for example, a complete telomere-to-telomere human X chromosome recently being assembled^8^.

The cost of long-read sequence data can still be prohibitive for species with larger genomes, and the critical target for average read length and read depth remains unclear. A full assessment of the impacts of varying sequence read length and depth on the contiguity and completeness of assemblies is therefore essential for informed allocation of finite resources. Here we conduct a comprehensive assembly experiment using subsets of a high-depth, long-read (PacBio) data set for the maize inbred line NC358 to evaluate critical inflection points of quality during the assembly of a complex, repeat-rich genome.

We sequenced the NC358 genome to 75x depth (based on a ~2.27 Gb genome size^9^) using the PacBio Sequel platform, which generated a raw read N50 of 21.2 kb (**Table 1; Table S1; Figure S1**). To identify an optimal assembly approach for this study, the complete raw data from NC358 and data from the B73 v4 genome assembly (68x depth)^10^ were each assembled using Falcon^11^, Canu^12^, and a hybrid approach in which Falcon was used for error correction and Canu was used for assembly. All assembled contigs were superscaffolded with a *de-novo* Bionano optical map (**Figure S2**), and pseudomolecules were constructed based on maize GoldenGate genetic markers^13^ and high-density maize pan-genome markers^14^ (Online Methods). The Falcon-Canu hybrid assemblies of both genomes showed consistently higher quality in terms of contig length, Bionano conflicts, Benchmarking Universal Single-Copy Orthologs (BUSCOs)^15^, and LTR Assembly Index (LAI)^7^ (**Table S2**), thus this method was used for all subsequent assemblies performed on subsets of the data.

**Table 1.**
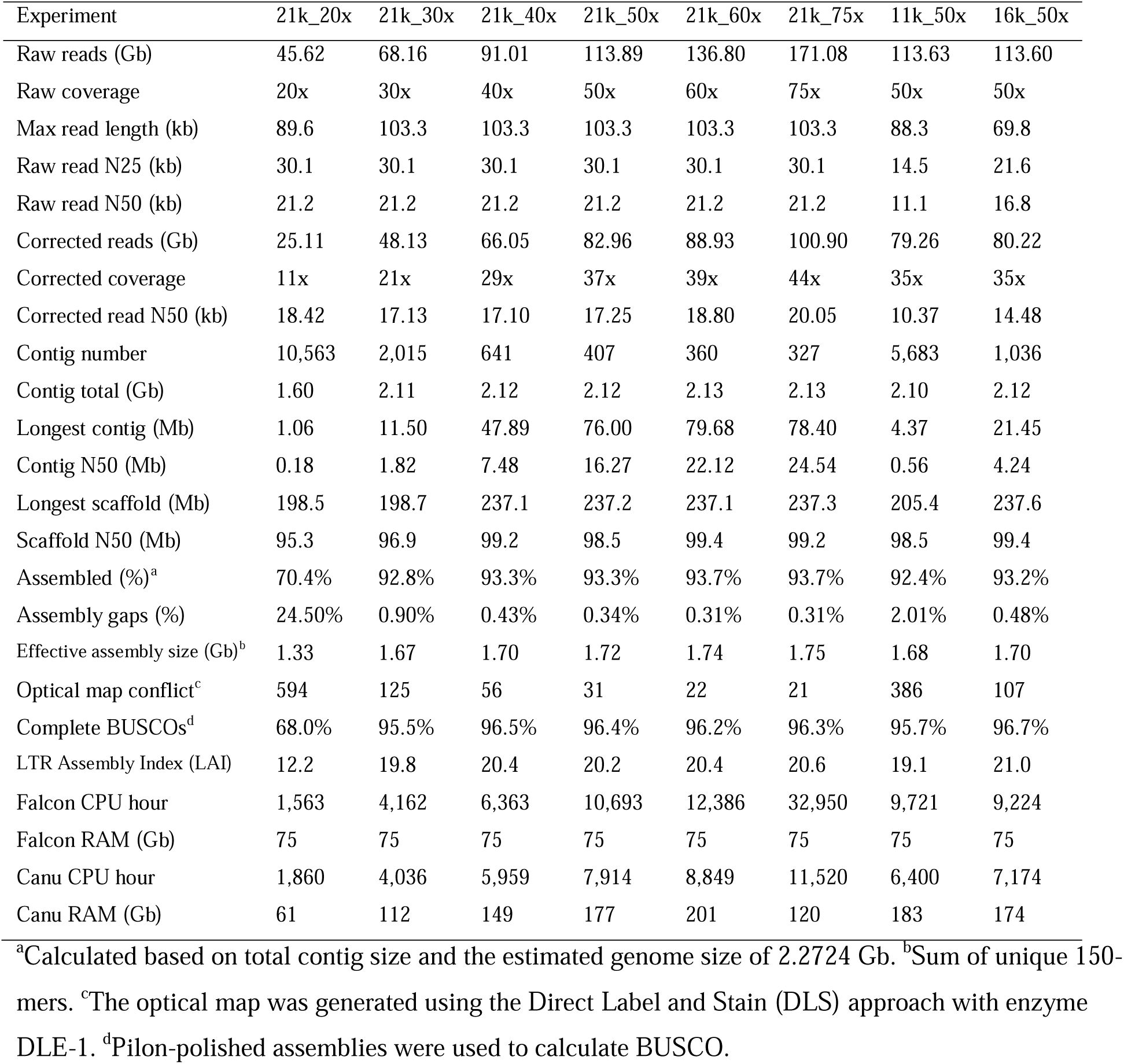
Summary statistics for NC358 assemblies.

| Experiment | 21k_20x | 21k_30x | 21k_40x | 21k_50x | 21k_60x | 21k_75x | 11k_50x | 16k_50x |
| --- | --- | --- | --- | --- | --- | --- | --- | --- |
| Raw reads (Gb) | 45.62 | 68.16 | 91.01 | 113.89 | 136.80 | 171.08 | 113.63 | 113.60 |
| Raw coverage | 20x | 30x | 40x | 50x | 60x | 75x | 50x | 50x |
| Max read length (kb) | 89.6 | 103.3 | 103.3 | 103.3 | 103.3 | 103.3 | 88.3 | 69.8 |
| Raw read N25 (kb) | 30.1 | 30.1 | 30.1 | 30.1 | 30.1 | 30.1 | 14.5 | 21.6 |
| Raw read N50 (kb) | 21.2 | 21.2 | 21.2 | 21.2 | 21.2 | 21.2 | 11.1 | 16.8 |
| Corrected reads (Gb) | 25.11 | 48.13 | 66.05 | 82.96 | 88.93 | 100.90 | 79.26 | 80.22 |
| Corrected coverage | 11x | 21x | 29x | 37x | 39x | 44x | 35x | 35x |
| Corrected read N50 (kb) | 18.42 | 17.13 | 17.10 | 17.25 | 18.80 | 20.05 | 10.37 | 14.48 |
| Contig number | 10,563 | 2,015 | 641 | 407 | 360 | 327 | 5,683 | 1,036 |
| Contig total (Gb) | 1.60 | 2.11 | 2.12 | 2.12 | 2.13 | 2.13 | 2.10 | 2.12 |
| Longest contig (Mb) | 1.06 | 11.50 | 47.89 | 76.00 | 79.68 | 78.40 | 4.37 | 21.45 |
| Contig N50 (Mb) | 0.18 | 1.82 | 7.48 | 16.27 | 22.12 | 24.54 | 0.56 | 4.24 |
| Longest scaffold (Mb) | 198.5 | 198.7 | 237.1 | 237.2 | 237.1 | 237.3 | 205.4 | 237.6 |
| Scaffold N50 (Mb) | 95.3 | 96.9 | 99.2 | 98.5 | 99.4 | 99.2 | 98.5 | 99.4 |
| Assembled (%)<sup>a</sup> | 70.4% | 92.8% | 93.3% | 93.3% | 93.7% | 93.7% | 92.4% | 93.2% |
| Assembly gaps (%) | 24.50% | 0.90% | 0.43% | 0.34% | 0.31% | 0.31% | 2.01% | 0.48% |
| Effective assembly size (Gb)<sup>b</sup> | 1.33 | 1.67 | 1.70 | 1.72 | 1.74 | 1.75 | 1.68 | 1.70 |
| Optical map conflict<sup>c</sup> | 594 | 125 | 56 | 31 | 22 | 21 | 386 | 107 |
| Complete BUSCOs<sup>d</sup> | 68.0% | 95.5% | 96.5% | 96.4% | 96.2% | 96.3% | 95.7% | 96.7% |
| LTR Assembly Index (LAI) | 12.2 | 19.8 | 20.4 | 20.2 | 20.4 | 20.6 | 19.1 | 21.0 |
| Falcon CPU hour | 1,563 | 4,162 | 6,363 | 10,693 | 12,386 | 32,950 | 9,721 | 9,224 |
| Falcon RAM (Gb) | 75 | 75 | 75 | 75 | 75 | 75 | 75 | 75 |
| Canu CPU hour | 1,860 | 4,036 | 5,959 | 7,914 | 8,849 | 11,520 | 6,400 | 7,174 |
| Canu RAM (Gb) | 61 | 112 | 149 | 177 | 201 | 120 | 183 | 174 |

<sup>a</sup>Calculated based on total contig size and the estimated genome size of 2.2724 Gb. <sup>b</sup>Sum of unique 150-mers. <sup>c</sup>The optical map was generated using the Direct Label and Stain (DLS) approach with enzyme DLE-1. <sup>d</sup>Pilon-polished assemblies were used to calculate BUSCO.

**Table 1.** Summary statistics for NC358 assemblies.
| Experiment | 21k_20x | 21k_30x | 21k_40x | 21k_50x | 21k_60x | 21k_75x | 11k_50x | 16k_50x |
| --- | --- | --- | --- | --- | --- | --- | --- | --- |
| Raw reads (Gb) | 45.62 | 68.16 | 91.01 | 113.89 | 136.80 | 171.08 | 113.63 | 113.60 |
| Raw coverage | 20x | 30x | 40x | 50x | 60x | 75x | 50x | 50x |
| Max read length (kb) | 89.6 | 103.3 | 103.3 | 103.3 | 103.3 | 103.3 | 88.3 | 69.8 |
| Raw read N25 (kb) | 30.1 | 30.1 | 30.1 | 30.1 | 30.1 | 30.1 | 14.5 | 21.6 |
| Raw read N50 (kb) | 21.2 | 21.2 | 21.2 | 21.2 | 21.2 | 21.2 | 11.1 | 16.8 |
| Corrected reads (Gb) | 25.11 | 48.13 | 66.05 | 82.96 | 88.93 | 100.90 | 79.26 | 80.22 |
| Corrected coverage | 11x | 21x | 29x | 37x | 39x | 44x | 35x | 35x |
| Corrected read N50 (kb) | 18.42 | 17.13 | 17.10 | 17.25 | 18.80 | 20.05 | 10.37 | 14.48 |
| Contig number | 10,563 | 2,015 | 641 | 407 | 360 | 327 | 5,683 | 1,036 |
| Contig total (Gb) | 1.60 | 2.11 | 2.12 | 2.12 | 2.13 | 2.13 | 2.10 | 2.12 |
| Longest contig (Mb) | 1.06 | 11.50 | 47.89 | 76.00 | 79.68 | 78.40 | 4.37 | 21.45 |
| Contig N50 (Mb) | 0.18 | 1.82 | 7.48 | 16.27 | 22.12 | 24.54 | 0.56 | 4.24 |
| Longest scaffold (Mb) | 198.5 | 198.7 | 237.1 | 237.2 | 237.1 | 237.3 | 205.4 | 237.6 |
| Scaffold N50 (Mb) | 95.3 | 96.9 | 99.2 | 98.5 | 99.4 | 99.2 | 98.5 | 99.4 |
| Assembled (%)<sup>a</sup> | 70.4% | 92.8% | 93.3% | 93.3% | 93.7% | 93.7% | 92.4% | 93.2% |
| Assembly gaps (%) | 24.50% | 0.90% | 0.43% | 0.34% | 0.31% | 0.31% | 2.01% | 0.48% |
| Effective assembly size (Gb)<sup>b</sup> | 1.33 | 1.67 | 1.70 | 1.72 | 1.74 | 1.75 | 1.68 | 1.70 |
| Optical map conflict<sup>c</sup> | 594 | 125 | 56 | 31 | 22 | 21 | 386 | 107 |
| Complete BUSCOs<sup>d</sup> | 68.0% | 95.5% | 96.5% | 96.4% | 96.2% | 96.3% | 95.7% | 96.7% |
| LTR Assembly Index (LAI) | 12.2 | 19.8 | 20.4 | 20.2 | 20.4 | 20.6 | 19.1 | 21.0 |
| Falcon CPU hour | 1,563 | 4,162 | 6,363 | 10,693 | 12,386 | 32,950 | 9,721 | 9,224 |
| Falcon RAM (Gb) | 75 | 75 | 75 | 75 | 75 | 75 | 75 | 75 |
| Canu CPU hour | 1,860 | 4,036 | 5,959 | 7,914 | 8,849 | 11,520 | 6,400 | 7,174 |
| Canu RAM (Gb) | 61 | 112 | 149 | 177 | 201 | 120 | 183 | 174 |

<sup>a</sup>Calculated based on total contig size and the estimated genome size of 2.2724 Gb. <sup>b</sup>Sum of unique 150-mers. <sup>c</sup>The optical map was generated using the Direct Label and Stain (DLS) approach with enzyme DLE-1. <sup>d</sup>Pilon-polished assemblies were used to calculate BUSCO.

Raw reads were downsampled from 75x to 60x, 50x, 40x, 30x, and 20x while maintaining a 21-kb raw-read N50 and to 50x depth with a raw-read N50 of 11 kb and 16 kb. These latter two data sets were generated to mirror read length distributions used in recent PacBio assemblies with similar genome sizes, including the human HG002 (ref. ^16^) and maize B73 v4 (ref. ^10^) genome assemblies (**Figure S3**). NC358 read subsets were error-corrected and assembled using the hybrid assembly approach described above (Online Methods; Supplementary Text). These processes were resource-intensive and were accelerated through cloud computing. The CPU time required for both Falcon error correction and Canu assembly increased substantially as read depth increased, while the required maximum memory was fairly similar (**Figure 1H; Table 1**).

**Figure 1.**
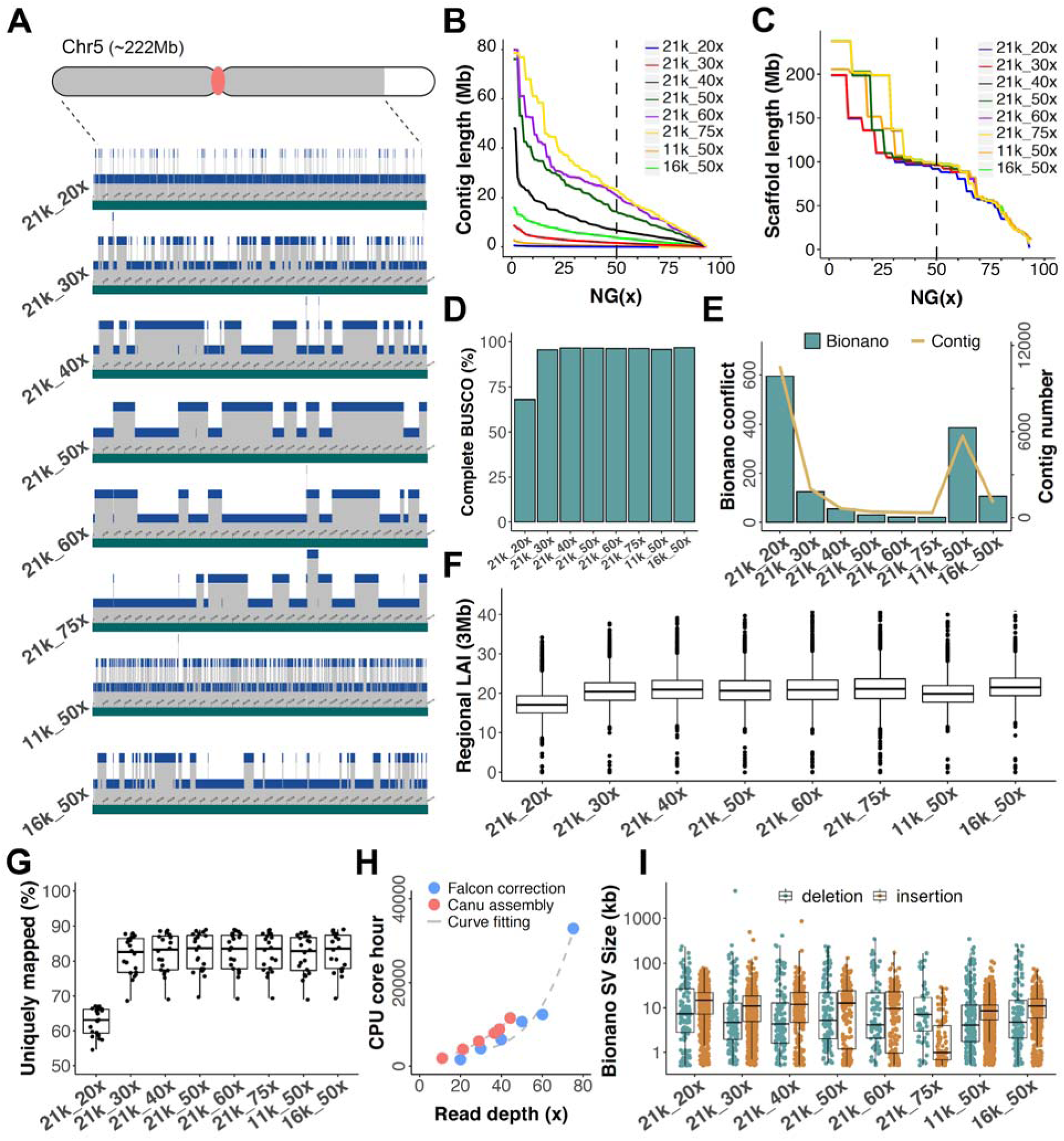
Assembly of NC358 using various read lengths and coverage. (A) Hybrid scaffolding using the Bionano optical map. A 199-Mb scaffold from chromosome 5 is shown. Grey areas on the chromosome cartoon represent the 199-Mb scaffold; the white area is the remaining 23-Mb scaffold in chromosome 5; the red dot is the centromere. Green tracts represent scaffolded sequences, and blue tracts show the contigs that comprise this scaffold with contigs jittered across three levels. (B) Contig NG(x). (C) Scaffold NG(x). (D) BUSCO. (E) The number of conflicts between Bionano contigs and sequence contigs and the number of contigs of each assembly. (F) Regional LAI values estimated based on 3-Mb windows with 300-kb steps. (G) Unique mapping rate of RNA-seq libraries. Each dot represents an RNA-seq library. (H) CPU core hours required for Falcon correction and Canu assembly. (I) Bionano optical map inconsistency. Deletions and insertions are cases where sequences are shorter or longer than the size estimated by the optical map, respectively.

Most assemblies had a total contig size covering >92% of the flow-cytometry estimated genome size of NC358 (2.27 Gb^9^), with the notable exception of the 21k_20x assembly (70.4% covered; **Table 1**). Contig length metrics were positively correlated with both read length and sequence coverage (**Figure 1B**), with the highest contig N50 (24.54 Mb) and the longest contig (79.68 Mb) observed in the 21k_75x and 21k_60x assembly, respectively (**Table 1**). A dramatic drop in quality was observed for both the lowest depth (21k_20x) and shortest sequence length (11k_50x) assemblies, where the number of contigs was 17x - 32x more than the complete 21k_75x dataset (**Table 1; Figure 1E)**.

For each assembly, superscaffolds were generated from the contigs using a common Bionano optical map. Even the most fragmented Falcon-Canu assembly could be scaffolded to high contiguity using this optical map due to the high density of labels in the map (**Figure 1A-C**). The resulting assemblies all had scaffold N50s at ~98 Mb (**Table 1**). In fact, chromosome 3 (~237 Mb) consisted of a single scaffold in five out of eight assemblies (**Table 1**). However, conflicts versus the Bionano map were much higher in the assemblies with 20x coverage and a raw-read N50 of 11 kb (**Table 1; Figure 1E**), suggesting assembly error increased with lower coverage and read length. Assemblies with shorter read length contained many more deletions relative to the optical map (**Figure 1I**), which may be due to the collapse of repetitive sequences. We did not observe a clear pattern between read length and deletion size (**Figure 1I**). Assembly misjoins were reduced with both longer reads and higher coverage, as shown by the relative number of insertions (**Figure 1I**).

For each of the assemblies, pseudomolecules were constructed using the GoldenGate and pan-genome genetic markers, which placed >99% of the total assembled bases into pseudomolecules (**Table S3; Figure S4**). The resulting NC358 pseudomolecules were highly syntenic across assemblies and to the B73 v4 genome (**Figure S5**).

We evaluated the completeness of gene-rich regions in each of the assemblies using BUSCO^15^. The percentage of complete BUSCO genes increased from 68.0% to 96.3% from the 21k_20x to the 21k_75x assembly (**Table 1; Figure 1D; Table S4**). Minimal improvement in BUSCO scores was achieved at depths higher than 30x (95.5% complete BUSCO genes), indicating this depth provides satisfactory gene space assembly.

To further evaluate the assembly of genic regions, we annotated gene models in the 21k_20x and the 21k_75x assemblies (Online Methods) and obtained a total of 28,275 and 39,578 genes, respectively (**Table S5**), with 92% of missing genes in the 21k_20x assembly falling within sequence gaps (**Table 1**). Exon and intron lengths of the annotated genes were similar across the assemblies (**Table S5**). Additionally, we sequenced RNA libraries from 10 tissues with two biological replicates (Online Methods). On average, 80% of reads in these libraries could be uniquely mapped to the various NC358 assemblies (**Figure 1G**). The 21k_20x assembly was a notable exception with only 63% of reads uniquely mapped (**Figure 1G; Figure S6**). We extracted the reads that did not map to the 21k_20x assembly and remapped them to the 21k_75x assembly, obtaining a unique-mapping rate of 36% (**Table S6**). These reads mapped to 3,184 genes in the 21k_75x assembly (**Table S7**). Of these 3,184 genes, 20% are present in the 21k_20x assembly but had assembly errors that prevented the RNA-seq reads from mapping, while the other 80% were within sequence gaps (**Table S7**).

In addition to metrics of gene completeness, we also examined each assembly for its ability to capture two notable maize tandem gene arrays, *Rp1-D*^17^ and *zein*^18^. The total length of these gene arrays was estimated at 536 kb and 62 kb in NC358 respectively based on the optical map. Both the *Rp1-D* and *zein* loci were completely assembled in all except for the 21k_20x assembly, where only 70% and 91% of the loci were assembled respectively (**Figure 2G; Table S8**).

**Figure 2.**
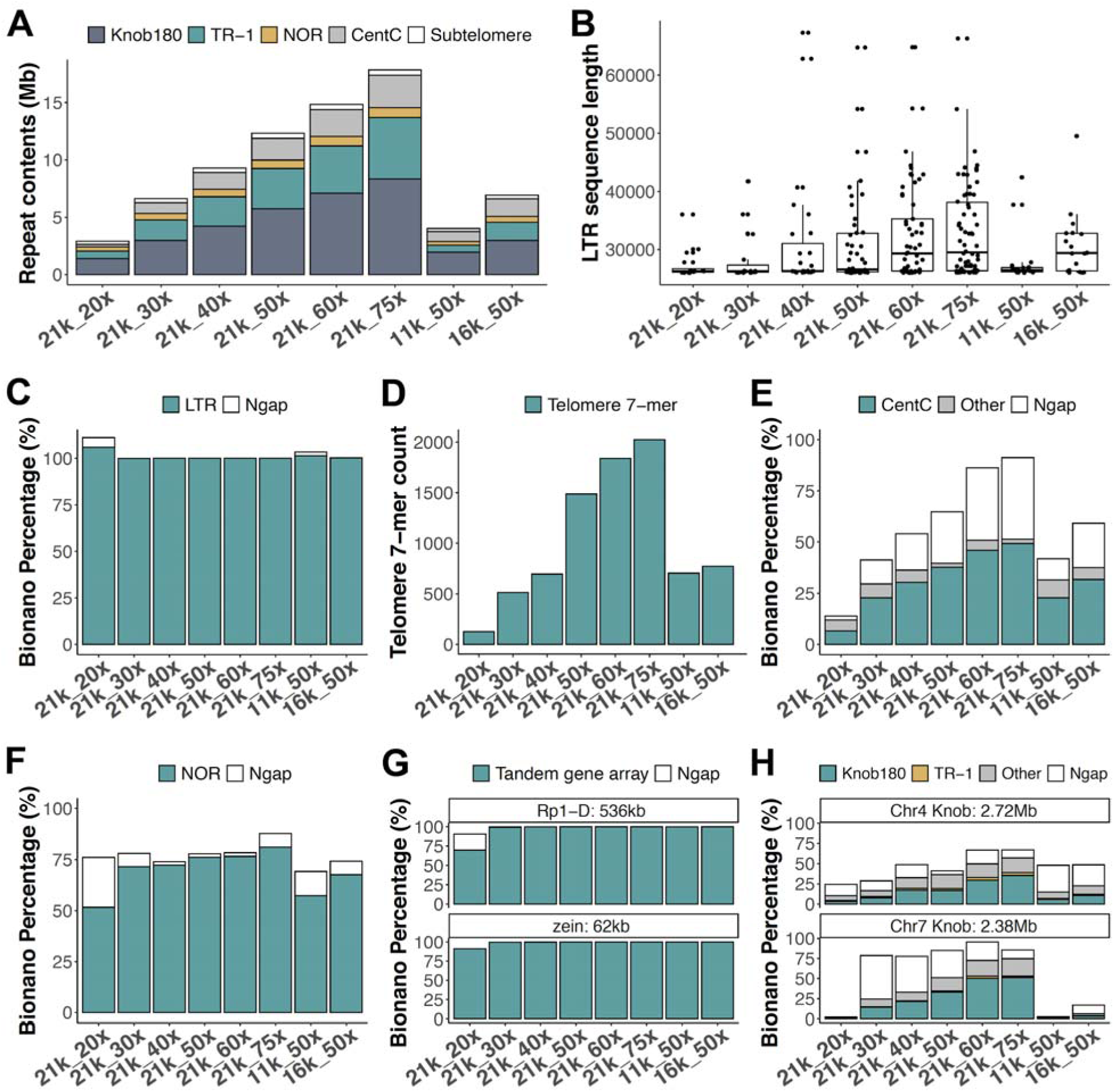
Assembly of repetitive components in the NC358 genome. (A) The assembled size of the 180-bp knob repeat, the knob TR-1 element, the chromosome 6 NOR region, CentC arrays, and subtelomere arrays in each of the NC358 assemblies. (B) Length distribution of LTR retrotransposons longer than 26 kb. Each dot represents an annotated sequence. (D) Telomere 7-mer counts in telomere regions of NC358 assemblies. Assembly of (C) LTR retrotransposons, (E) CentC arrays, (F) the chromosome 6 NOR region, (G) the *Rp1-D* and *zein* tandem gene arrays, and (H) two example knobs in each of the NC358 assemblies. The NC358 Bionano optical map was used to estimate the size of these components. Ngap, estimated gap size.

The completeness of transposon-rich regions of the genome was assessed through the assembly index of LTR retrotransposons, called LAI^7^. A higher LAI score is indicative of a more complete assembly in TE-rich regions. The 21k_20x assembly had a substantially lower LAI score compared to other assemblies (LAI = 12.2; **Table 1**). As sequence depth increased a substantial improvement in LAI was observed, while the effect of sequence length on LAI was minimal (**Figure 1F**). This is likely due to the fact that the length of LTR retrotransposons is approximately 10 kb on average (**Figure S7**), which could be spanned by even the 11 kb reads. The assemblies that were generated from ≥40x genomic depth achieved “gold” quality (LAI ≥ 20 (ref. ^7^)) (**Table 1; Figure 1F**), which was comparable to the B73 v4 genome and much higher than many previously published maize genome assemblies generated with short-read data (**Figure S8**).

The insertion time of each LTR retrotransposon can be dated based on sequence divergence between terminal repeats^7^. We identified 36% fewer intact LTR retrotransposons in the highly fragmented 21k_20x assembly (**Figure S9**), and significantly older LTR elements in the 11k_50x assembly (*p* < 10^−5^, Tukey’s test), suggesting fragmentation of assemblies could bias conclusions of transposon studies. LTR retrotransposons shorter than 26 kb were assembled well across the assemblies (**Figure S10; Figure S11**). However, a substantial effect of longer reads and higher depth was observed in the assembly of LTR sequences longer than 26 kb (**Figure 2B**). We examined the assemblies of the longest LTR sequence clusters using the Bionano optical map and found most assemblies contained no gaps and were virtually complete (**Figure 2C**), with the notable exception that the 11k_50x, 16k_50x, and 21k_20x assemblies, which contained large gaps in one of the LTR clusters (**Table S9**). We also inspected the *bz* locus^19^, which has highly nested transposon insertions and an estimated size of 303.5 kb in NC358. The *bz* locus was well assembled in all but the 21k_20x assembly, in which only 56.3% of the sequence was included (**Table S10**). In summary, with ≥40x of sequence coverage, long-read sequencing and assembly can traverse most transposon-rich genomic regions including relatively long LTR sequences, though with shorter reads (*i.e.*, read N50 of 11 kb - 16 kb) this sequencing depth may not be sufficient.

The assembly of non-TE tandem repeat space was also evaluated, including telomeres (7-bp repeats), subtelomeres (300 - 1300-bp repeats), CentC arrays (156-bp repeats), nucleolus organizer region (NOR, ~11 kb repeats), and the two major knob repeats (mixture of 180-bp and 350-bp repeats) (**Figure 2A; Table S11**). The effects of sequence read depth and sequence read length were far more pronounced across many of these tandemly duplicated portions of the genome (**Figure 2A**).

Telomeres are characterized by 7-bp tandem repeats at the end of each chromosome. Our results showed a substantial increase in the assembled length of telomere sequence with the increase of both read length and sequence coverage (**Figure 2D; Table S12**). However, a precise estimate of telomere length was not possible with our optical map due to the lack of Bionano DLE-1 sites in these highly repetitive regions. Using the full dataset (21k_75x), only 10 of 20 telomere-subtelomere combined regions were assembled to >90% of the Bionano estimated size (**Table S13**), suggesting even longer reads and higher coverage are required for the full assembly of these regions.

The centromere is one of the most repetitive regions of many species’ genomes including maize. We characterized NC358 centromeres based on CentC arrays^20^ which are abundant in functional centromeric regions^21^. Even with the full dataset (21k_75x), only half of CentC arrays were assembled (**Figure 2E; Figure S12; Table S14**). Hybrid scaffolded assemblies with sequence coverage ≥60x yielded a better approximation to the Bionano estimated size, even though these regions largely consisted of gaps (**Figure 2E**). Although assembled sequences were not significantly increased, higher sequence depth resulted in better anchoring of sequences with the Bionano optical map. Only three centromeres, which contained a mixture of CentC arrays, transposons, and intergenic sequences, could be traversed by Bionano DLE-1 labeling due to having a comparatively higher content of low-copy sequence^21^. The size of the remaining centromeres was likely underestimated (**Figure S13**), and further improvements in scaffolding technology are required to traversing these regions.

The NOR is enriched with ribosomal DNA (rDNA) and spans approximately 9 Mb on chromosome 6 of NC358 (**Table S15**). Longer read length improved the assembly of this region, but substantial differences were not observed with coverage ≥30x (**Figure 2F**). Approximately 72% of the NOR was included in the 21k_30x assembly and this improved by just 9% to 81% in the 21k_75x assembly (**Table S15; Figure 2F**).

Finally, maize knobs are heterochromatic regions consisting of 180-bp (knob180) and 350-bp (TR-1) repeats^22^. We used the Bionano optical map to assess the assembly of two knobs that together spanned a total of 5 Mb. With longer reads and higher coverage, more knob sequences were assembled, with 6.5% of the two knobs present in the 21k_20x assembly and up to 65% in the 21k_75x assembly (**Table S16**; **Figure 2H**).

Recent innovations in long-read and scaffolding technology have made highly contiguous assembly possible across a wide range of species. We have documented how both the completeness and contiguity of assemblies improve with increasing depth and read length. The biological aims of an investigation must be considered when determining the level of investment in depth of sequence. With long-read sequencing, the low-copy gene space (including tandem gene arrays) can be well assembled with as low as 30x genomic coverage across a range of read lengths. Complete characterization of transposable elements in complex genomes such as maize will require a greater depth of sequence (~40x) and should employ library preparation protocols that maximize read-length N50. Finally, complete assembly of highly repetitive genomic features such as heterochromatic knobs, telomeres, and centromeres will require substantially more data. In fact, complete assembly of these latter highly repetitive sequences will likely require innovations beyond current sequencing technology.

## ONLINE METHODS

### Sample preparation

Seeds for the maize NC358 inbred line were obtained from GRIN Global (seed stock ID Ames 27175), grown, and self-pollinated at Iowa State University in 2017. A total of 144 seedlings derived from a single selfed ear were grown in the greenhouse. Leaf tissues from the seedlings at the Vegetative 2 (V2) growth stage were sampled after a 48-hour dark treatment to reduce carbohydrates. A total of 35g of tissue was harvested and flash-frozen. Tissue was sent to the Arizona Genomics Institute (AGI) for high molecular weight DNA isolation using a CTAB protocol^23^.

### Illumina and PacBio Sequencing

Pacific BioSciences long-read data for NC358 were generated at AGI using the Sequel platform. Libraries were prepared using the manufacturer’s suggested protocol (https://www.pacb.com/). The raw reads that were generated covered the genome at an estimated 75-fold depth (75x) with a read-length N50 of 21,166 bp. Reads from each SMRT cell were inspected and quality metrics were calculated using SequelQC^24^. After validating the PSR (polymerase to subread ratio) and ZOR (ZMW occupancy ratio) were satisfactory, all subreads were used for subsequent steps.

Paired-end Illumina data for NC358 were generated at the Georgia Genomics and Bioinformatics Core (GGBC) from the same DNA extraction as was used for the long-read sequencing. Quality control of DNA was conducted using Qubit and Fragment Analyzer to determine the concentration and size distribution of the DNA. The library was constructed using the KAPA Hyper Prep Kit (Cat# KK8504). During library preparation, DNA was fragmented by acoustic shearing with Covaris before end repair and A-tailing. Barcoded adaptors were ligated to DNA fragments to form the final sequencing library. Libraries were purified and cleaned with SPRI beads before being amplified with PCR. Final libraries underwent another bead cleanup before being evaluated by Qubit, qPCR (KAPA Library Quantification Kit Cat# KK4854), and Fragment Analyzer. The final pool undergoing Illumina’s Dilute and Denature Libraries protocol was diluted to 2.2 pM for loading onto the sequencer and then sequenced with 1% PhiX by volume. Libraries were sequenced on the NextSeq 500 instrument using PE150 cycles. The demultiplexing was done on Illumina’s BaseSpace.

PacBio SMRT subreads for the maize inbred line B73 (sequenced to 68x depth) were retrieved from the NCBI SRA database with accession ID SRX1472849 (ref. ^10^). PacBio SMRT subreads for the human HG002 sample (sequenced to 147x depth) were retrieved with accession IDs SRX1033793 and SRX1033794 (ref. ^16^).

### Downsampling raw sequence

The 75x SMRT Sequel raw data from maize NC358 was downsampled to 60x, 50x, 40x, 30x, and 20x data using seqtk (v1.2) (https://github.com/lh3/seqtk). Downsampling was performed as serial titration, in which each dataset was the superset of the next smaller dataset, and was sampled to have similar length distributions (**Figure S3**). The N50 of the downsampled raw data were almost identical to the N50 of the full 75x data (**Table 1**).

### Shifting read length distribution of raw sequence

Two more NC358 datasets were downsampled and trimmed from the original 75x SMRT dataset to match the read length distribution of the maize B73 data^10^ and the human HG002 data^16^, which had read N50 lengths of ~16 kb and ~11 kb, respectively (**Figure S3**). To do this, first, the read lengths of the maize B73 and human HG002 data were each sorted in descending order. For each read length value, all raw reads from NC358 that were longer than said value were randomly sampled without replacement and clipped to have matched read length. The unused clipped part of the read was put back in the pool for further use with short read length. This distribution-shifting approach was chosen to achieve a realistic distribution of read length rather than trimming all reads by fixed lengths. These datasets were labeled as “16k”, and “11k” based on their N50 of raw data of 16,765, and 11,092, respectively.

### RNA tissue sampling and sequencing

Samples from 10 tissues throughout development were collected to generate expression evidence for gene annotation. Two biological replicates were collected for each tissue type, and each replicate consisted of three individual plants. The tissues that were sampled were: 1) primary root at six days after planting; 2) shoot and coleoptile at six days after planting; 3) base of the 10^th^ leaf at the Vegetative 11 (V11) growth stage; 4) middle of the 10^th^ leaf at the V11 growth stage; 5) tip of the 10^th^ leaf at the V11 growth stage; 6) meiotic tassel at the Vegetative 18 (V18) growth stage; 7) immature ear at the V18 growth stage; 8) anthers at the Reproductive 1 (R1) growth stage; 9) endosperm at 16 days after pollination; and 10) embryo at 16 days after pollination. Tissue from developmental stage V11 and older were taken from field-grown plants while all younger tissue samples were taken from greenhouse-grown plants. For the endosperm and embryo samples, tissue from 50 kernels per plant (150 total per biological replicate) were sampled. Greenhouse-grown plants were planted in Metro-Mix300 (Sun Gro Horticulture) with no additional fertilizer and grown under greenhouse conditions (27°C/24°C day/night and 16h/8h light/dark) at the University of Minnesota Plant Growth Facilities. Field grown plants were planted at the Minnesota Agricultural Experiment Station located in Saint Paul, MN with 30-inch row spacing at ~52,000 plants per hectare. RNA was extracted using the Qiagen RNeasy plant mini kit following the manufacturer’s suggested protocol.

The quality of the total RNA was assessed by Bioanalyzer or Fragment analyzer to determine RNA concentration and integrity. The sample concentration was normalized in 25 uL of nuclease-free H_2_O before library preparation. Libraries were prepared using KAPA’s stranded mRNA-seq kit with halved reaction volumes. During library preparations, mRNA was selected using oligo-dT beads, the RNA was fragmented, and cDNA was generated using random hexamer priming. Single or dual indices were ligated depending on the desired level of multiplexing. The number of cycles for library PCR was determined based on kit recommendations for the amount of total RNA used during library preparation. Libraries were quality control checked using Qubit or plate reader, depending on the number of samples in the batch for library concentration, and fragment analyzer for the size distribution of the library. The pooling of samples was based on qPCR. The pooled libraries were then checked by Qubit, Fragment Analyzer, and qPCR.

RNA libraries were prepared for sequencing on Illumina instruments using Illumina’s Dilute and Denature protocol. Pooled libraries were diluted to 4 nM, then denatured using NaOH. The denatured library was further diluted to 2.2 pM, and PhiX was added at 1% of the library volume. RNA pools were sequenced on a NextSeq 550 to generate 75 bp pair-end reads. On average, 24.5 million pair-end reads were generated per replicate per tissue type, for a total of 489 million reads across all samples. Data were demultiplexed and trimmed of adapter and barcode sequences on BaseSpace (**Figure S14)**.

### Bionano data generation

The DNA extraction was performed using the Bionano Prep™ Plant Tissue DNA Isolation Kit according to a modified version of the Plant Tissue DNA Isolation Base Protocol. Approximately 0.5g leaf tissue was collected from young etiolated seedlings germinated in soil-free conditions and grown in the dark for approximately two weeks after germination. Freshly-cut leaves were treated with a 2% formaldehyde fixing solution and then washed, cut into small pieces and homogenized using a Qiagen TissueRuptor probe. Free nuclei were concentrated by centrifugation at 2000 xg, washed, isolated by gradient centrifugation and embedded into a low-melting-point agarose plug. After proteinase K and RNase A treatments, the agarose plug was washed four times in Wash Buffer and five times in TE (Tris and EDTA) buffer. Finally, purified ultra-high molecular weight nuclear DNA (uHMW nDNA) was recovered by melting the plug, digesting it with agarase and subjecting the resulting sample to drop dialysis against TE.

The Bionano Saphyr platform, in combination with the Direct Label and Stain (DLS) process, was used to generate chromosome-level sequence scaffolds and validate PacBio sequence contigs. Direct labeling was performed using the Direct Labeling and Staining Kit (Bionano Genomics Catalog 80005) according to the manufacturer’s recommendations, with some modifications^25^. In total, 1 ug uHMW nDNA was incubated for 2:20 h at 37 °C, followed by 20 min at 70 °C in the presence of DLE-1 Enzyme, DL-Green and DLE-1 Buffer. Following proteinase K digestion and cleanup of the unincorporated DL-Green label, the labeled DNA was combined with Flow Buffer, DTT, and incubated overnight at 4 °C. DNA was quantified and stained by adding Bionano DNA Stain to a final concentration of 1 microliter per 0.1 microgram of final DNA. The labeled sample was loaded onto a Bionano chip flow cell and molecules separated, imaged and digitized in a Bionano Genomics Saphyr System and server according to the manufacturer’s recommendations (https://bionanogenomics.com/support-page/saphyr-system/).

Data visualization, processing, DLS map assembly, and hybrid scaffold construction were all performed using the Bionano Genomics software Access, Solve, and Tools. A filtered subset of 1,282,746 molecules (353,596 Mb total length) with a minimum size of 150 kb and a maximum size of 3 Mb were assembled without pre-assembly using the non-haplotype parameters with no CMPR cut and without extend-split.

### Genome assembly

To determine the assembly approach to apply to each of the datasets, three different methods were first tested on the complete dataset, including Falcon only, Canu only, and a Falcon-Canu hybrid approach. We also downloaded raw PacBio sequencing data for the B73 v4 genome for comparison of the different approaches with a second data set.

The Falcon genome assemblies were performed using the falcon_kit pipeline v0.7 (ref. ^11^) with some modifications. TANmask and REPmask were not used due to their extensive masking for the maize genome. Error correction for raw reads was performed on the longest 50x coverage, with the average read correction rate set to 75% (-e 0.75) and local alignments for at least 3000 bp (-l 3000). The usage of -l 3000 instead of -l 2500 was done because of the omitted repeat masking, which works better for highly repetitive genome species like maize. A minimum of two reads and a maximum of 200 reads were used for error corrections (--min_cov 2 --max_n_read 200). For sequence assembly, the exact matching k-mers between two reads was set to 24 bp (-k 24) with read correction rate as 95% (-e 0.95) and local alignments at least 1000 bp (-l 1000). The longest 20x coverage reads were used for assembly with a minimum coverage of two and maximum coverage of 80 (--min_cov 2 --max_cov 80). Full parameter sets are included in the supplementary text.

For Canu read correction and assembly, Canu v1.7 (ref. ^12^) was used. K-mers more frequent than 500 were not used to seed overlaps (ovlMerThreshold=500). The genome size of 2,272,400,000 bp and 2,500,000,000 bp for NC358 and B73, respectively, were used in this study^9^. Other parameters were used as default. Due to a bug in the Canu v1.7 program, truncations of large contigs would occur during the consensus process (https://github.com/marbl/canu/releases/tag/v1.8). Because the program was not expecting the superlong contigs that were being generated for our NC358 assemblies, we found a total of nine large contigs that suffered from consensus truncations. To fix these truncation gaps, consensus-free contigs were generated using Canu v1.7 (cnsConsensus=quick), then blastn was used to search for 5-kb boundaries of truncation gaps in consensus-free assemblies. Truncated sequences were retrieved and patched to the truncated contigs.

For the Falcon-Canu hybrid approach, the error correction was performed by Falcon, and the trimming and assembly were performed by Canu using the versions and parameters described above. All the assemblies were performed on the DNAnexus cloud platform. CPU core hour and maximum memory usage were recorded every 10 minutes for each Falcon error correction and Canu assembly job. For Falcon error correction of the 21k datasets, the CPU core hour (y) could be predicted by raw read depth (m) with y = 20603100000 + (3136.685 - 20603100000)/(1 + (m/1932.377)^4.148144). For Canu assembly of the 21k datasets, the CPU core hour (y) could be predicted by corrected read depth (n) with y = 6438752000 + (1284.689 - 6438752000)/(1 + (n/56334.74)^1.872455). These curves were fit using the https://mycurvefit.com/ website and plotted in R.

We evaluated these assembly approaches using both maize NC358 and B73. For both inbred lines, a similar assembly size was generated by each of the approaches. However, the Falcon-Canu hybrid approach yielded the longest contig length (78.4 Mb and 19.7 Mb, respectively), the highest contig NG50 (23.0 Mb and 3.0 Mb, respectively), and the lowest number of assembly errors based on Bionano conflict cuts (21 and 64, respectively; **Table S1**). The gene space completeness evaluated using Benchmarking Universal Single-Copy Orthologs (BUSCOs)^15^ and the repeat space continuity evaluated using the LTR Assembly Index (LAI) (vbeta3.2)^7^ were similar between the Canu and the hybrid approach and higher than those assemblies that were created using the Falcon assembler (**Table S1**). This was likely due to the consensus approach used at the end of the Canu program, which was missing in the Falcon program. Due to the consistently high quality of the assemblies generated from the hybrid approach, we used this approach to assemble each of the NC358 datasets with varying sequence depth and read length. Full parameter sets are included in the supplementary text.

### Genome polishing

Two polishing approaches were tested on the 21k_75x assembly. The first was done using Arrow with PacBio raw reads (75x coverage). Read mapping to the assembly was done using BLASR^26^ with default parameters (--minMatch 12 --bestn 10 --minPctSimilarity 70.0 --refineConcordantAlignments). The Arrow tool in the SMRT Link (v5.1.0) software package was then applied to correct for sequencing errors with default parameters. A second approach for polishing was done using Pilon with Illumina pair-end reads (30.7x coverage). Read mapping to the assembly was done using Minimap2 (v2.16)^27^ with the short read option (-ax sr). Pilon (v1.23-0)^28^ was then applied to correct for sequencing errors including SNPs and small indels (--fix bases) on sites with a minimum depth of 10 and a minimum mapping quality of 30 (--mindepth 10 --minmq 30).

With both approaches, minimal differences were observed in the contiguity statistics (**Table S2**) or the repeat content for the 21k_75x assembly (**Figure S15**), and it is expected that this minimal impact would be observed across all of the NC358 assemblies. A more substantial difference in BUSCO scores were observed with both the Arrow-polished and the Pilon-polished 21k_75x assemblies (**Table S2**). Because the polishing had a substantial impact on this metric, the other NC358 assemblies were also polished using Pilon with the same parameter settings and similar improvement of BUSCO scores were observed (**Table 1; Table S4**).

### Generation of pseudomolecules

Hybrid scaffolds for the assemblies were generated with Bionano Direct Label and Stain data using Bionano Solve (v3.2.1_04122018). Overlaps of contigs within Bionano map space were resolved by placing 13 bp of Ns (13N gaps) at the overlap site. In addition to arranging contigs into scaffolds, the hybrid scaffold was also used to detect misassembly and to assess completeness of the assembled genome and repeat elements.

The pseudomolecules were constructed from the hybrid scaffolds using ALLMAPS (v0.8.12)^29^. Both pan-genome anchor markers^14^ and GoldenGate markers^13^ were used with equal weights for ordering and orientating the scaffolds. For pan-genome anchor markers, data were downloaded from the CyVerse Data Commons (http://datacommons.cyverse.org/browse/iplant/home/shared/panzea/genotypes/GBS/v27/Lu_2015_NatCommun_panGenomeAnchors20150219.txt.gz) and a bed file with 50 bp upstream and downstream of the B73 v3 coordinates were generated. A text file with marker name and predicted distance was also constructed from the same file. The extracted markers were mapped to HiSat2 (v2.1.0)^30^ indexed assemblies of NC358 by disabling splicing (--no-spliced-alignment) and forcing global alignment (--end-to-end). Very high read and reference gap open and extension penalties (--rdg 10000,10000 and --rfg 10000,10000) were also used to ensure full-length mapping of marker sequence. The final alignment was then filtered for mapping quality of greater than 30 and tag XM:0 (unique mapping) to retain only high-quality uniquely mapped marker sequences. The mapped markers were merged with the predicted distance information to generate a CSV input file for ALLMAPS. Only scaffolds with more than 20 uniquely mapped markers, with a maximum of 100 markers per scaffold, were used for pseudomolecule construction.

The GoldenGate markers were downloaded from MaizeGDB (https://www.maizegdb.org/data_center/map?id=1203673). For the markers with coordinates, 50 bp flanking regions were extracted from the B73 v4 genome. For markers without coordinates, marker sequences were used as-is, and those missing both coordinates and sequences were discarded. Mapping of the markers was done similar to the method described above for the pan-genome anchor markers, with all uniquely mapped markers retained. The genetic distance information for these markers was converted to a CSV file before using it in ALLMAPS. ALLMAPS was run with default options, and the pseudomolecules were finalized after inspecting the marker placement plot and the scaffold directions. Synteny dotplots were generated using the scaffolds as well as pseudomolecule assemblies against the B73 genome by following the ISUgenomics Bioinformatics Workbook (https://bioinformaticsworkbook.org/dataWrangling/genome-dotplots.html)^31^. Briefly, the repeats were masked using RepeatMasker (v4.0.9)^32^ and the Maize TE Consortium (MTEC) curated library^33^. RepeatMasker was configured to use the NCBI engine (rmblastn) with a quick search option (-q) and GFF as a preferred output. The repeat-masked genomes were then aligned using Minimap2 (v2.2)^27^ and set to break at 5% divergence (-x asm5). The paf files were filtered to eliminate alignments less than 1 kb and dotplots were generated using the R package dotPlotly (https://github.com/tpoorten/dotPlotly).

### Gene annotation and RNA-seq mapping

The MAKER-P pipeline^34^ was used to annotate protein-coding genes for Pilon-polished NC358 21k_20x and 21k_75x genome assemblies. The baseline evidence used in annotating the B73 v4 genome^10^ was applied. Before gene annotation, the MTEC curated TE library^33^ and RepeatMasker was used to mask repetitive sequences. For gene prediction, we used Augustus^35^ and FGENESH^36^ (http://www.softberry.com/berry.phtml) with training sets based on maize and monocots, respectively. To identify genes that were missing in the 21k_20x assembly, total coding sequences (CDS) from the 21k_75x annotation was masked by total CDS from the 21k_20x annotation using Repeatmasker (-div 2 - cutoff 1000 -q -no_is -norna -nolow). The 21k_75x CDS that were masked less than 20% were determined missing in the 21k_20x annotation. These missing CDS were blast against the 21k_20x assembly and those that had less than 20% similarity were also determined to be missing in the 21k_20x assembly.

A total of 20 RNA-seq libraries were sequenced from NC358 tissue samples. Each library was sequenced to 21.9x ± 0.7x coverage with a mapping rate of 86.4% ± 1.0% to the B73 v4 using STAR (v2.5.2b)^37^ (**Figure S16; Table S17**). To benchmark the gene space assembly, STAR (v2.5.2b)^37^ was used to map the RNA-seq reads against the Pilon-polished NC358 assemblies. Unmapped reads from the 21k_20x assembly were extracted using SAMtools^38^ and remapped to the 21k_75x assembly with STAR. Genes with read coverage ≥20% were extracted using BEDtools^39^, and blast against the 21k_20x assembly for the identification of full-length copies. The NC358 TE library (see next section for details on library generation) was used to identify TE fragments in genes with aligned reads (**Table S7**). In addition, TEsorter (v1.1.4)^40^ (https://github.com/zhangrengang/TEsorter) was used to identify TE-related protein domains in genes with default parameters (**Table S7**).

### Assessment of genome assembly quality

The quality of the different NC358 assemblies was assessed on the unpolished assemblies unless noted. For continuity, N50, NG50, NG(x), the number of contigs, and maximum contig length were estimated. NG(x) values were the length of the contig at the top x percent of the estimated genome size (2.2724 Gb) consisting of the longest contigs. NG50 is a commonly used case of NG(x) values. NG(x) values were calculated using GenomeQC (https://github.com/HuffordLab/GenomeQC)^41^. The gene space completeness was estimated using BUSCO (v3.0.2)^15^ with the Embryophyta odb9 dataset (n = 1,440) and BLAST (v2.6)^42^, Augustus (v3.3)^35^, EMBOSS (v6.6.0)^43^, and HMMER (v3.1b2)^44^.

The repeat space contiguity was accessed using the LTR Assembly Index (LAI) (vbeta3.2)^7^. To annotate LTR retrotransposons, LTR_retriever (v2.6)^45^ was used to identify intact LTR retrotransposons and construct LTR libraries for each NC358 assembly with default parameters. To generate a high-quality LTR library for NC358, assembly-specific LTR libraries were aggregated and masked by the MTEC curated LTR library using RepeatMasker (v4.0.7)^32^. Library sequences masked over 90% were removed and redundant sequences were also removed using utility scripts (cleanup_tandem.pl and cleanup_tandem.pl) from the EDTA package^46^. Non-redundant NC358-specific LTR sequences were added to the MTEC curated LTR library to form the final LTR library for NC358. The final library was then used to mask the 21k_75x assembly for the estimation of total LTR content. The total LTR content of 76.34% and LTR identity of 94.854% was used to estimate LAI values of all NC358 assemblies (-totLTR 76.34 -iden 94.854). The LAI of the other maize line genomes, including PH207 (GeneBank Accession: GCA_002237485.1)^47^, CML247 (GeneBank Accession: GCA_002682915.2)^14^, Mo17 (From Xin *et al*. (2013)^48^ and GeneBank Accession: GCA_003185045.1 (ref. ^49^)), W22 (GeneBank Accession: GCA_001644905.2)^50^, and B73 v4 (GeneBank Accession: GCA_000005005.6)^10^ were also evaluated for context.

Effective assembly size, which is the length of the uniquely mappable sequences of an assembly, was estimated using unique 150-mers in each sequence assembly and quantified using Jellyfish (v2.0)^51^ with default parameters.

### Misassembly identification with optical maps

The Bionano optical mapping was used as an orthogonal method to identify misassemblies in genomes. Bionano *de novo* assembled optical maps were aligned to the sequence pseudomolecules to characterize structural inconsistencies using the structural variant calling pipeline of BionanoSolve 3.4. Default parameters were employed from the nonhaplotype_noES_DLE file. Homozygous calls with a confidence of 0.1, a size of 500 bp, and non-overlaps with gap regions were regarded as insertions and deletions in sequence assemblies.

### Assembly quality evaluation in repeat space

The coordinates of CentC arrays, knob180, TR-1 knobs, and NOR in the assemblies were identified by blasting CentC, knob180, TR-1 knob consensus sequences^21^, and the rDNA intergenic spacer (AF013103.1) against each assembly. An individual repeat array was defined as clusters of repetitive sequences that had less than 100 kb interspace between repeated elements. The level of repeats and gaps were then quantified in each defined repeat array. Respective sizes of each repeat array in the Bionano maps were estimated using the Bionano labels closest to the start and end coordinates in the assemblies.

To identify the telomere-subtelomere boundaries of the NC358 assemblies, seven maize subtelomere repeat sequences were downloaded from NCBI (EU253568.1, S46927.1, S46926.1, S46925.1, CL569186.1, AF020266.1, and AF020265.1) and used as queries to blast against the NC358 21k_75x assembly. Subtelomere boundaries were first identified at the start and end of chromosomes where blast hits were clustering then cross-checked with subtelomere-specific Fluorescence in situ hybridization (FISH) data^52^. The blast results were concordant with FISH results, showing the beginning of chromosomes 7, 8, 9, and 10 lack subtelomeres (**Table S13**). Telomeres were defined as the distance between the boundary of subtelomeres to the end of pseudomolecules of the 21k_75x assembly, which were used as the basis for estimating the telomere size and count of the telomeric repeat sequences (5’-TTTAGGG-3’ and 5’-CCCTAAA-3’ in reverse complementation) in all other NC358 assemblies.

To identify the *bz* locus in the NC358 assemblies, the sequence of the maize W22 *bz* locus was first downloaded from NCBI (EU338354.1)^19^. The starting and ending 2 kb of the W22 *bz* locus were used to blast against the NC358 21k_75x assembly and the longest matches on chromosome 9 were used as the location of the *bz* locus in the NC358 21k_75x assembly. The obtained NC358 *bz* locus is 289,103 bp in length (chr9:11625031..11914133), which is 50 kb longer than that of the W22 *bz* locus (238,141 bp). Similarly, the 2-kb flanking sequences of the NC358 21k_75x *bz* locus were used to locate the *bz* locus coordinates in the other NC358 assemblies.

The *zein* sequence was downloaded from NCBI (AF031569.1) and the *Rp1-D* from MaizeGDB (AC152495.1_FG002). The same method as described for the *bz* locus was used to identify coordinates in the NC358 assemblies based on blast results using 2-kb flanking sequences.

## Data availability

PacBio and Illumina sequencing reads for the NC358 line used in this study are available with EBI Biosample ID ERSXXXXXXX. All code developed for this study is available on GitHub: https://github.com/HuffordLab/Maize_NC358.

## ACKNOWLEDGMENTS

This work was supported by NSF Plant Genome Research Program grant IOS-1744001 to RKD, DW, and MBH and grant IOS-1546727 to CNH, USDA ARS 5030-21000-068-00D to MBH and MW, and USDA ARS 58-8062-2100-044 to DW. BPW, SK, and AMP were supported by the Intramural Research Program of the National Human Genome Research Institute. We wish to acknowledge Jonathan Gent for helpful discussion on repeat space analyses.

## AUTOHR CONTRIBUTIONS

RKD, CNH, MBH, and DW conceived the study. AF, CSC, SO, and AS assembled the genomes. SO, JL, KMC, AF, AS, JS, VL, NM, AMG, XW, CSC, DEH, SP, SS, KF, MW, BPW, SK, AMP, and BH collected data and conducted the analyses. SO, JL, AF, AS, VL, RKD, CNH, MBH, DW wrote the manuscript. All authors read and approved the final manuscript.

## COMPETING INTERESTS STATEMENT

The authors declare that they have no competing interests.

## Notes

https://github.com/HuffordLab/Maize_NC358

