## Supplementary material for "Effect of Sequence Depth and Length in Long-read Assembly of the Maize Inbred NC358": Figure S (from another version of this preprint: version 1, 2019-11-29)

### SUPPLEMENTARY TEXT

#### Parameter for FALCON genome assembly

FALCON only assembly for 75-fold NC358:

pa_HPCdaligner_option = -k14 -e0.75 -s100 -l3000 -h240 -w8 -H14154

ovlp_HPCdaligner_option = -k24 -e.95 -s100 -l1000 -h600 -H20149

pa_DBsplit_option = -x500 -s400

ovlp_DBsplit_option = -x500 -s400

falcon_sense_option = --min_idt 0.70 --min_cov 2 --max_n_read 200

overlap_filtering_setting = --max_diff 40 --max_cov 80 --min_cov 2 --min_len 500

Canu only assembly for 75-fold NC358:

ovlMerThreshold=500; genome_size=2272400000; input_type=-pacbio-raw

FALCON-Canu hybrid assembly for 60-fold NC358:

FALCON

pa_HPCdaligner_option = -k14 -e0.75 -s100 -l3000 -h240 -w8 -H10310

pa_DBsplit_option = -x500 -s400

falcon_sense_option = --min_idt 0.70 --min_cov 2 --max_n_read 200

CANU

ovlMerThreshold=500; genome_size=2272400000; input_type=-pacbio-corrected

FALCON-Canu hybrid assembly for 50, 40, 30, 20-fold NC358:

FALCON

pa_HPCdaligner_option = -k14 -e0.75 -s100 -l3000 -h240 -w8 -H3000

pa_DBsplit_option = -x500 -s400

falcon_sense_option = --min_idt 0.70 --min_cov 2 --max_n_read 200

CANU

ovlMerThreshold=500; genome_size=2272400000; input_type=-pacbio-corrected

FALCON-Canu hybrid assembly for 50-fold NC358 with shift distribution:

FALCON

pa_HPCdaligner_option = -k14 -e0.75 -s100 -l3000 -h240 -w8 -H3000

pa_DBsplit_option = -x500 -s400

falcon_sense_option = --min_idt 0.70 --min_cov 2 --max_n_read 200

CANU

ovlMerThreshold=500; genome_size=2272400000; input_type=-pacbio-corrected

FALCON only assembly for 68-fold B73:

pa_HPCdaligner_option = -k14 -e0.75 -s100 -l3000 -h240 -w8 -H9898

ovlp_HPCdaligner_option = -k29 -e.95 -s100 -l4800 -h600 -H12360

pa_DBsplit_option = -x500 -s400

ovlp_DBsplit_option = -x500 -s400

falcon_sense_option = --min_idt 0.70 --min_cov 2 --max_n_read 200

overlap_filtering_setting = --max_diff 40 --max_cov 80 --min_cov 2 --min_len 500

Canu only assembly for 68-fold B73:

ovlMerThreshold=500; genome_size=2500000000; input_type=-pacbio-raw

FALCON-Canu hybrid assembly for 68-fold B73:

FALCON

pa_HPCdaligner_option = -k14 -e0.75 -s100 -l3000 -h240 -w8 -H9898

pa_DBsplit_option = -x500 -s400

falcon_sense_option = --min_idt 0.70 --min_cov 2 --max_n_read 200

CANU

ovlMerThreshold=500; genome_size=2272400000; input_type=-pacbio-corrected

### SUPPLEMENTARY FIGURES

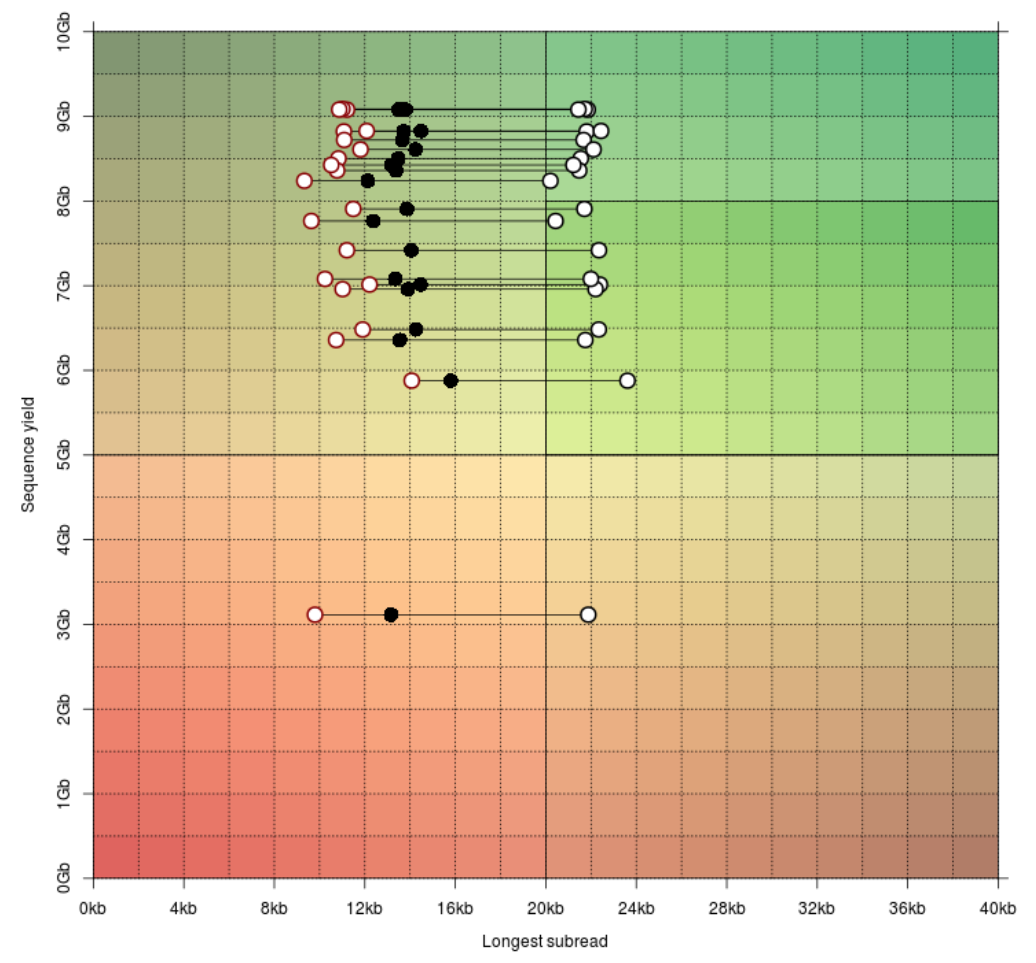

Figure S1. Statistics for NC358 subreads derived from 21 SMRT cells. X-axis, subread length. For a continuous long read (CLR) with multiple passes, only the longest subread was used. Y-axis, sequence yield. Each line represents a SMRT cell with black dots, white dots with red outlines, and white dots with black outlines represent subread mean, median, and N50 lengths, respectively.

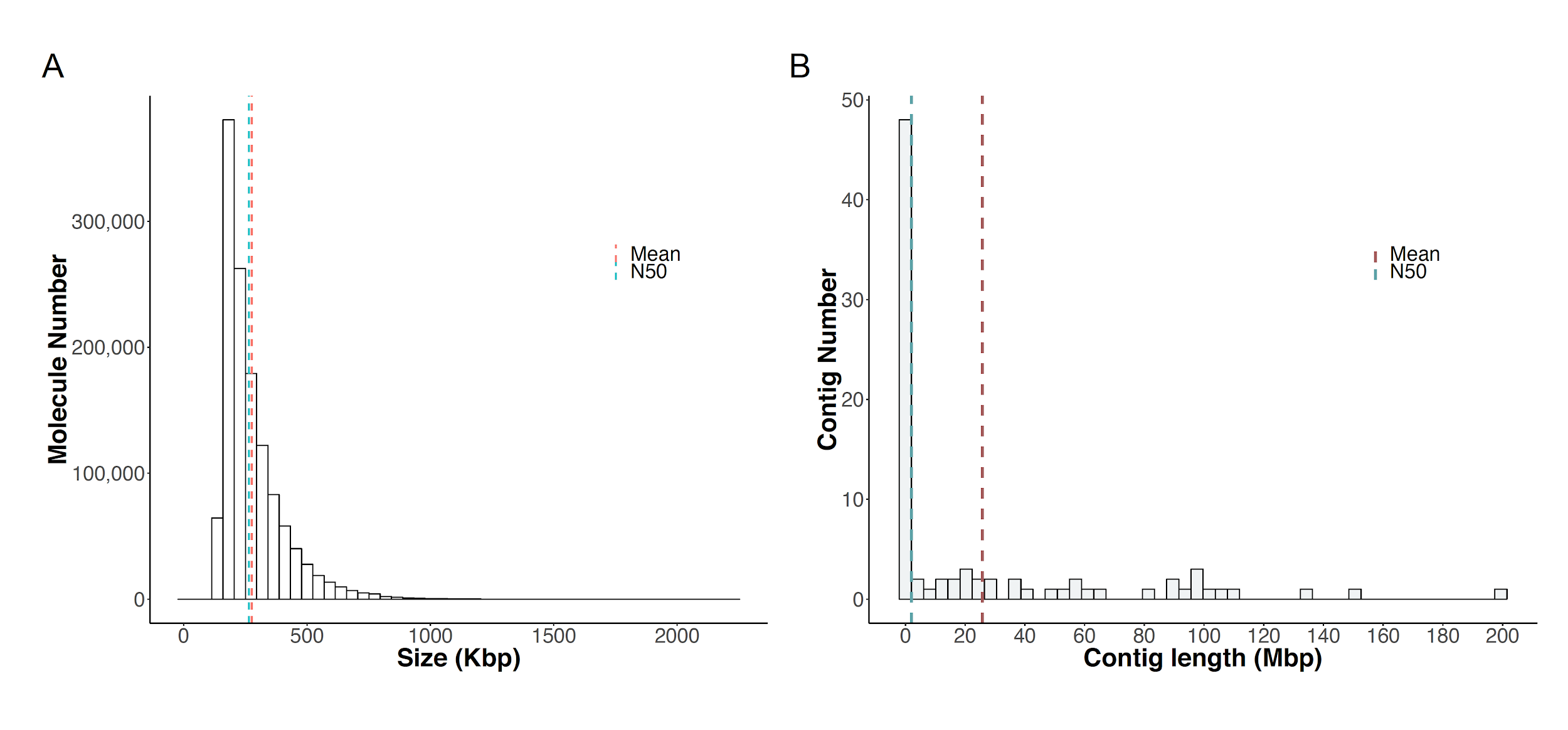

Figure S2. Summary of the NC358 Bionano optical map. (A) BNX molecule length distribution. (B) Bionano contig length distribution.

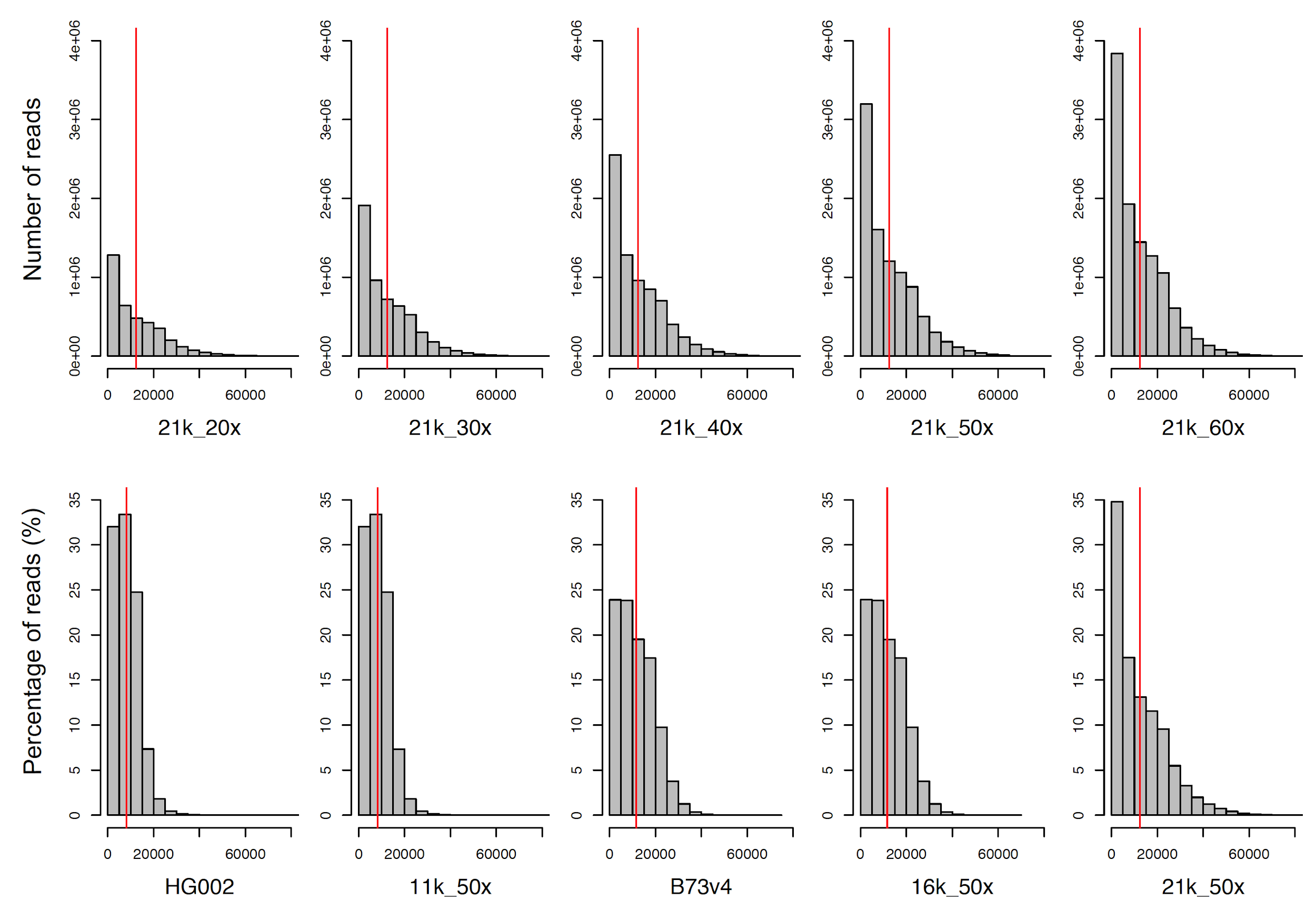

Figure S3. Raw read length distribution of different datasets. X-axes, read length in bp. Top row, raw read count was presented. Bottom row, read density was presented. Red vertical line, mean read length of the dataset.

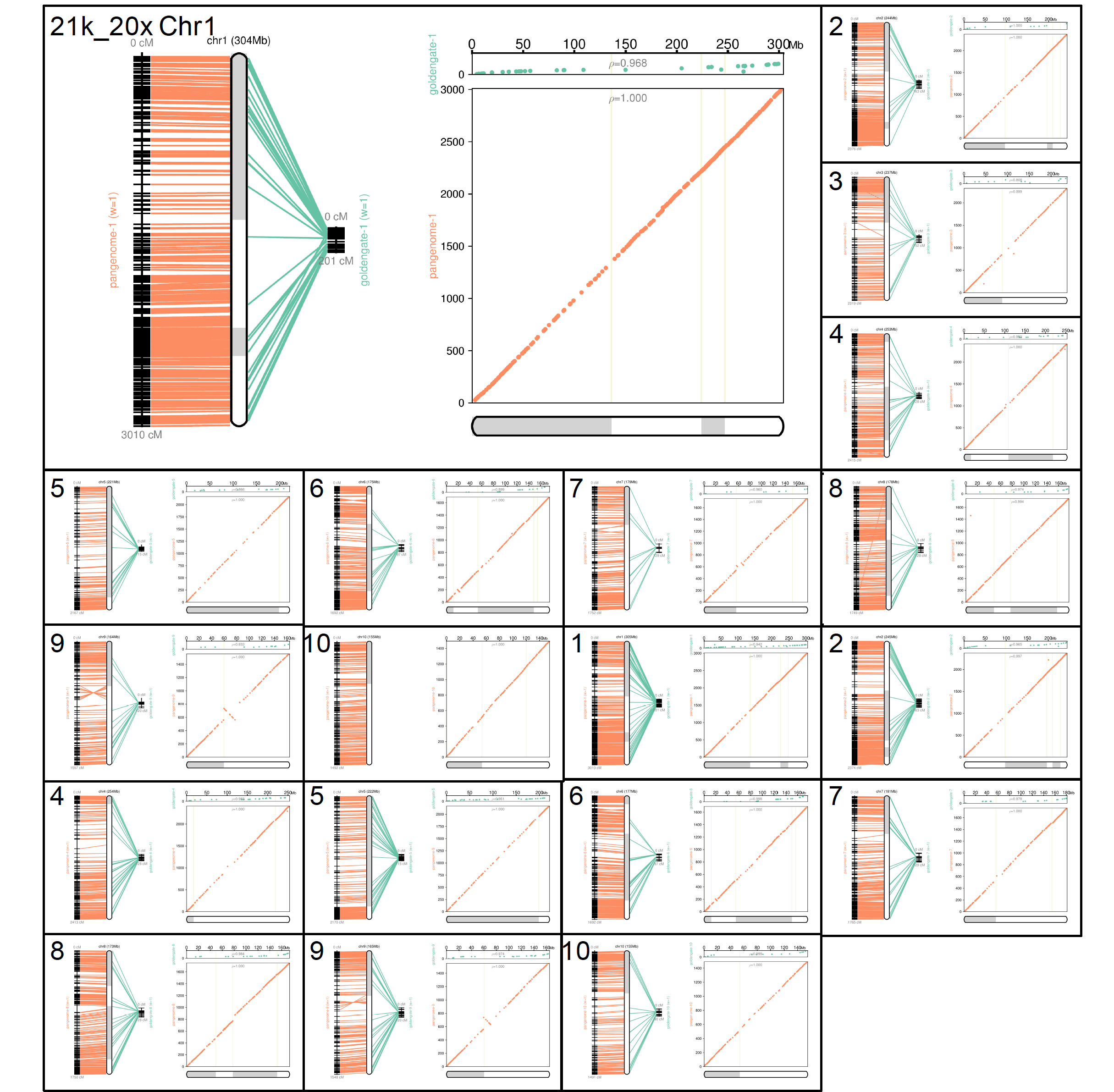

Figure S4. Construction of pseudomolecules using both GoldenGate and Pangenome markers. The NC358 21k_20x and 21k_75x are shown on the top and bottom, respectively, with each chromosome indicated on a single subplot. The 21k_75x chromosome 3 consists of a single scaffold, thus no pseudomolecule construction is needed and therefore not shown in the figure. Orange lines and dots, pangenome markers mapped to the pseudomolecule; Green lines and dots, GoldenGate markers mapped to the pseudomolecule; Grey and white regions, different superscaffolds of the pseudomolecule. Plots for the rest of the NC358 assemblies are available here: <https://github.com/HuffordLab/Maize_NC358/tree/master/AGP/AGP_plots>.

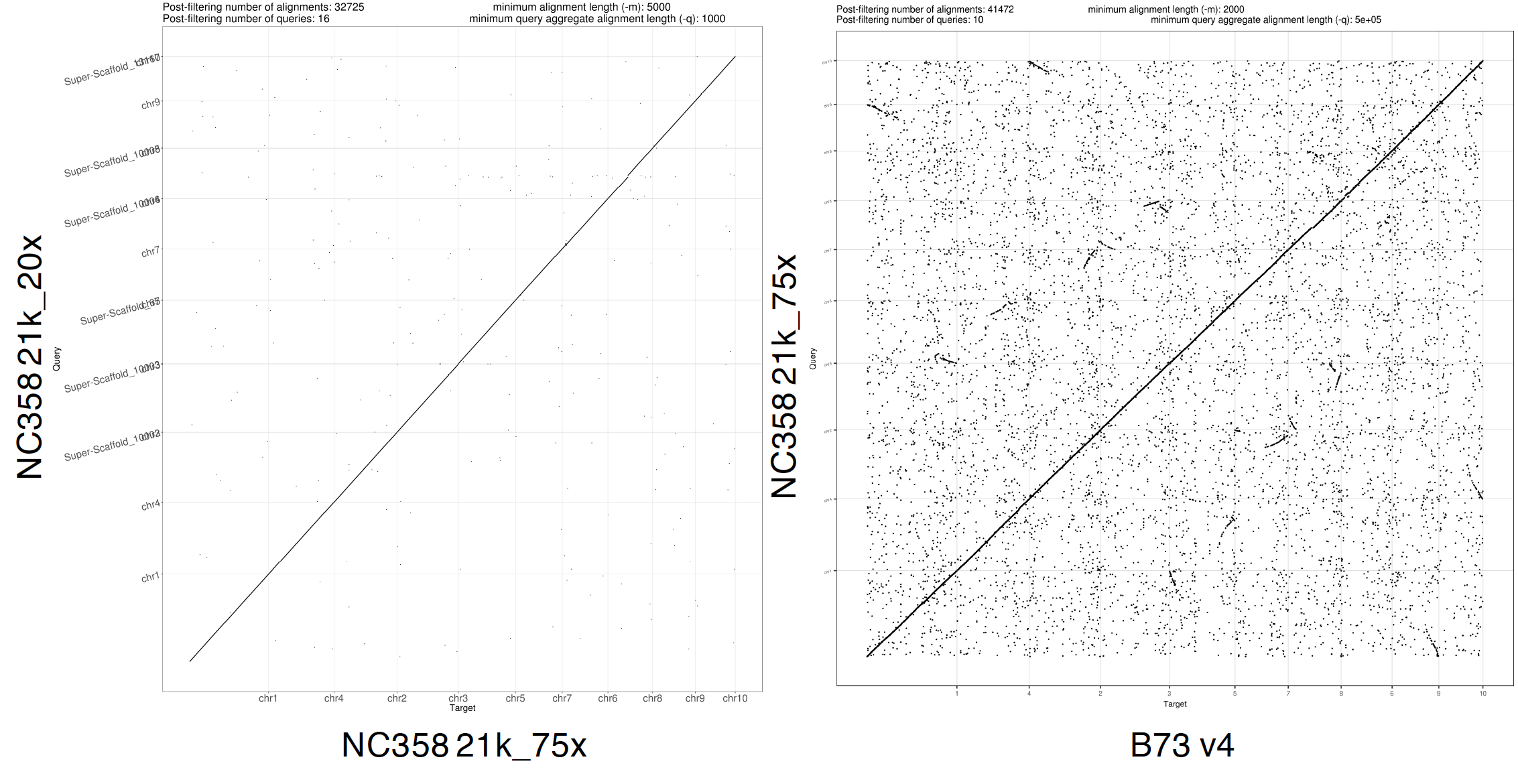

Figure S5. Synteny between NC358 21k_20x, NC358 21k_75x, and B73 v4 genomes.

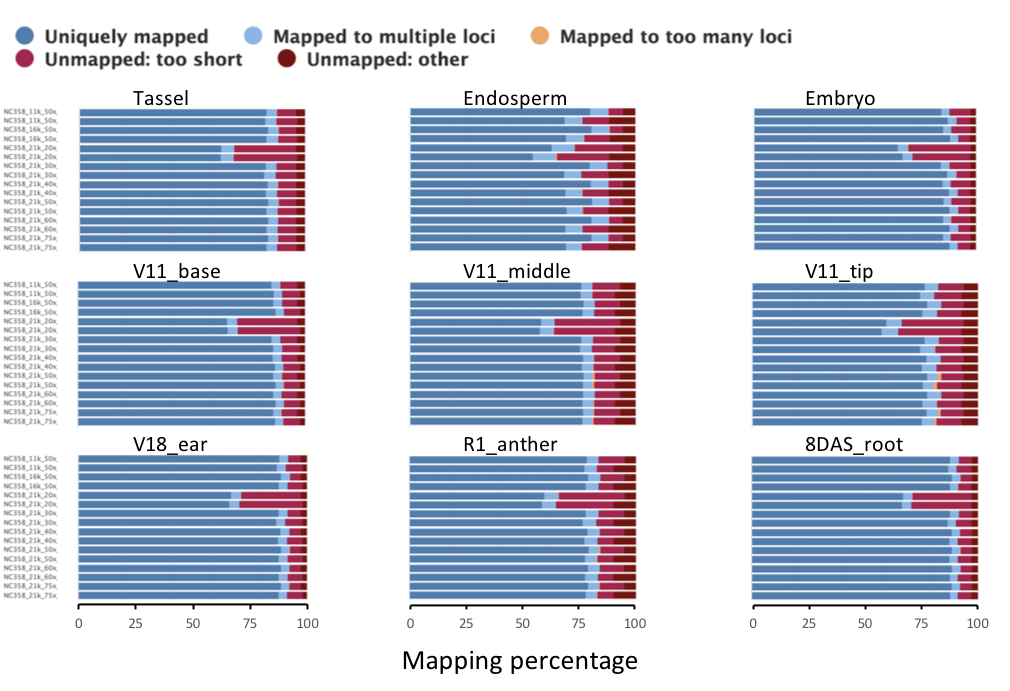

Figure S6. RNA-seq mapping of nine tissues to various NC358 assemblies. Each replicate (total two for each tissue) was mapped independently. The 21k_20x assembly showed significantly low RNA-seq mapping rates.

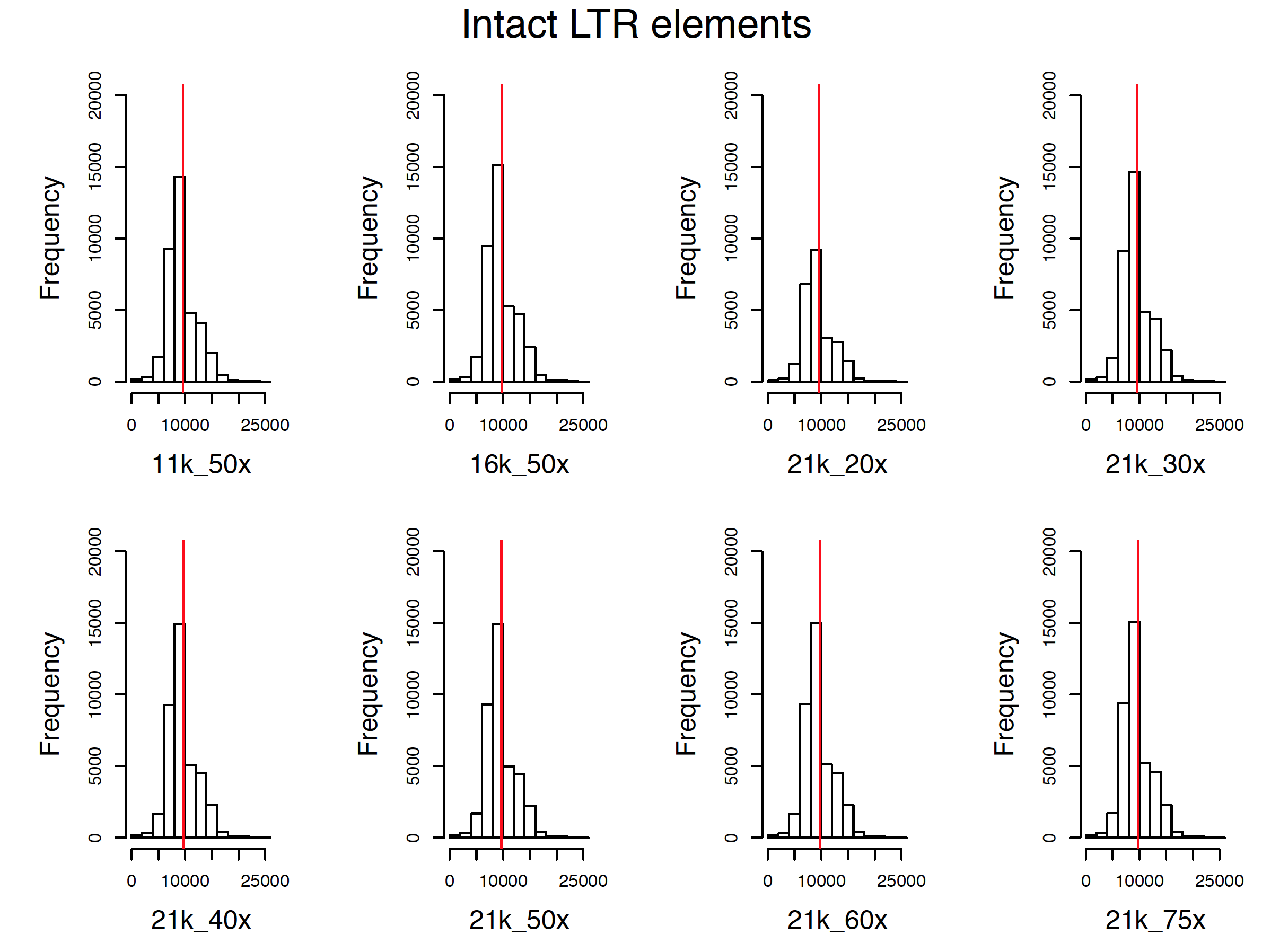

Figure S7. Length distribution of intact LTR retrotransposons in NC358 assemblies. X-axes, length of LTR retrotransposons. Y-axes, count of LTR retrotransposons. Red lines, mean length of LTR retrotransposons.

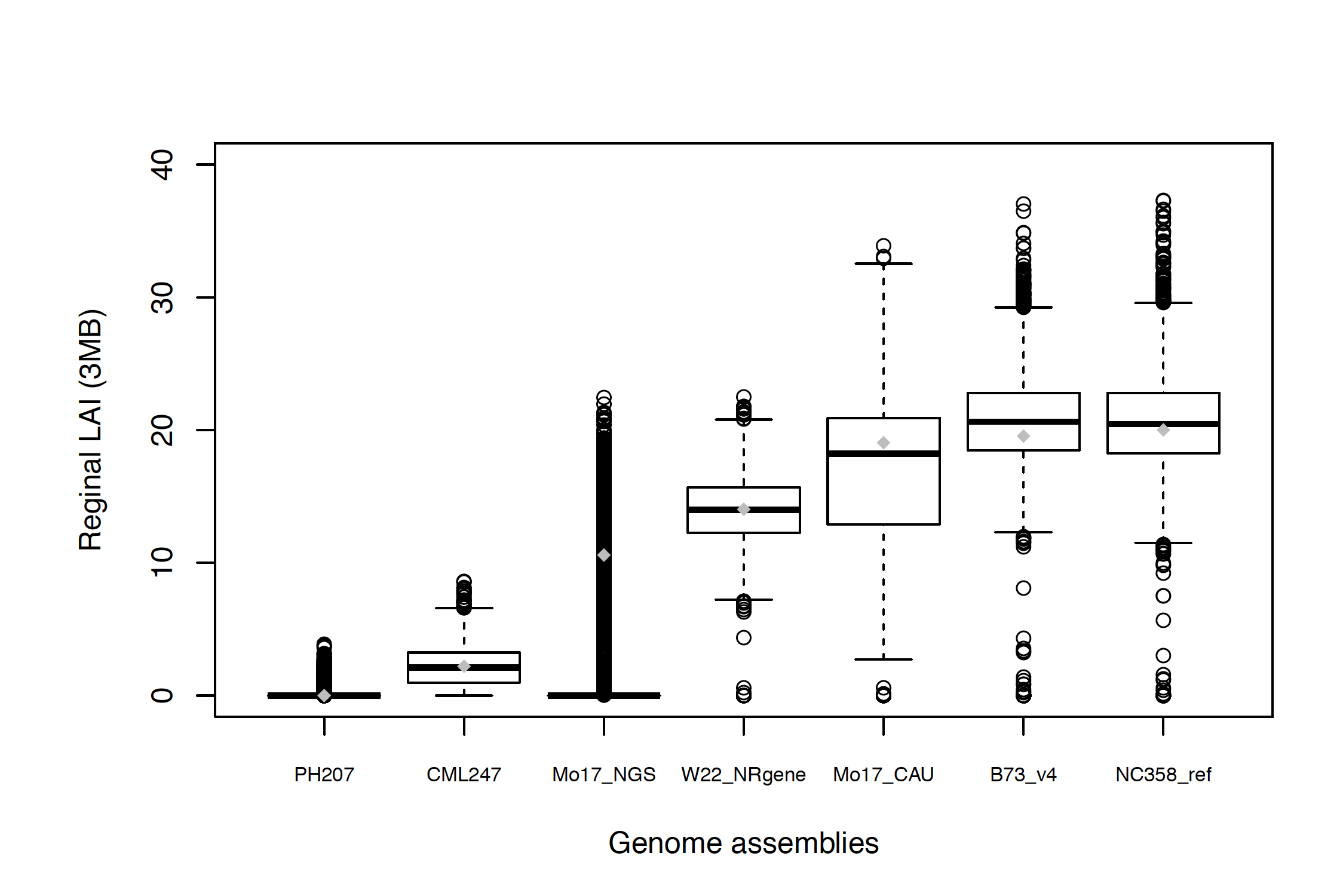

Figure S8. LTR Assembly Index (LAI) of NC358 and published genomes. Grey dot, whole-genome LAI.

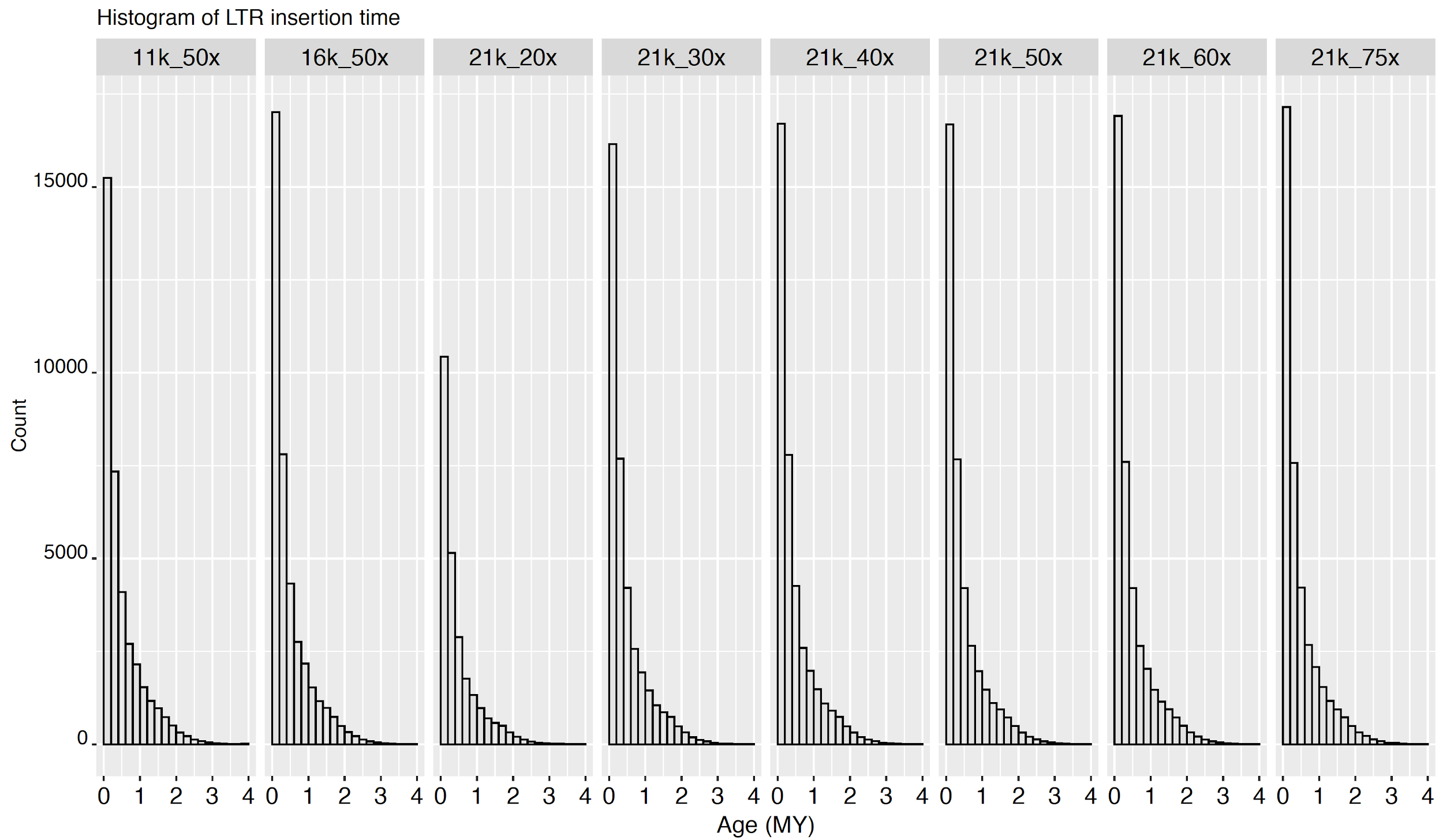

Figure S9. Distribution of insertion time of intact LTR retrotransposons identified in NC358 assemblies.

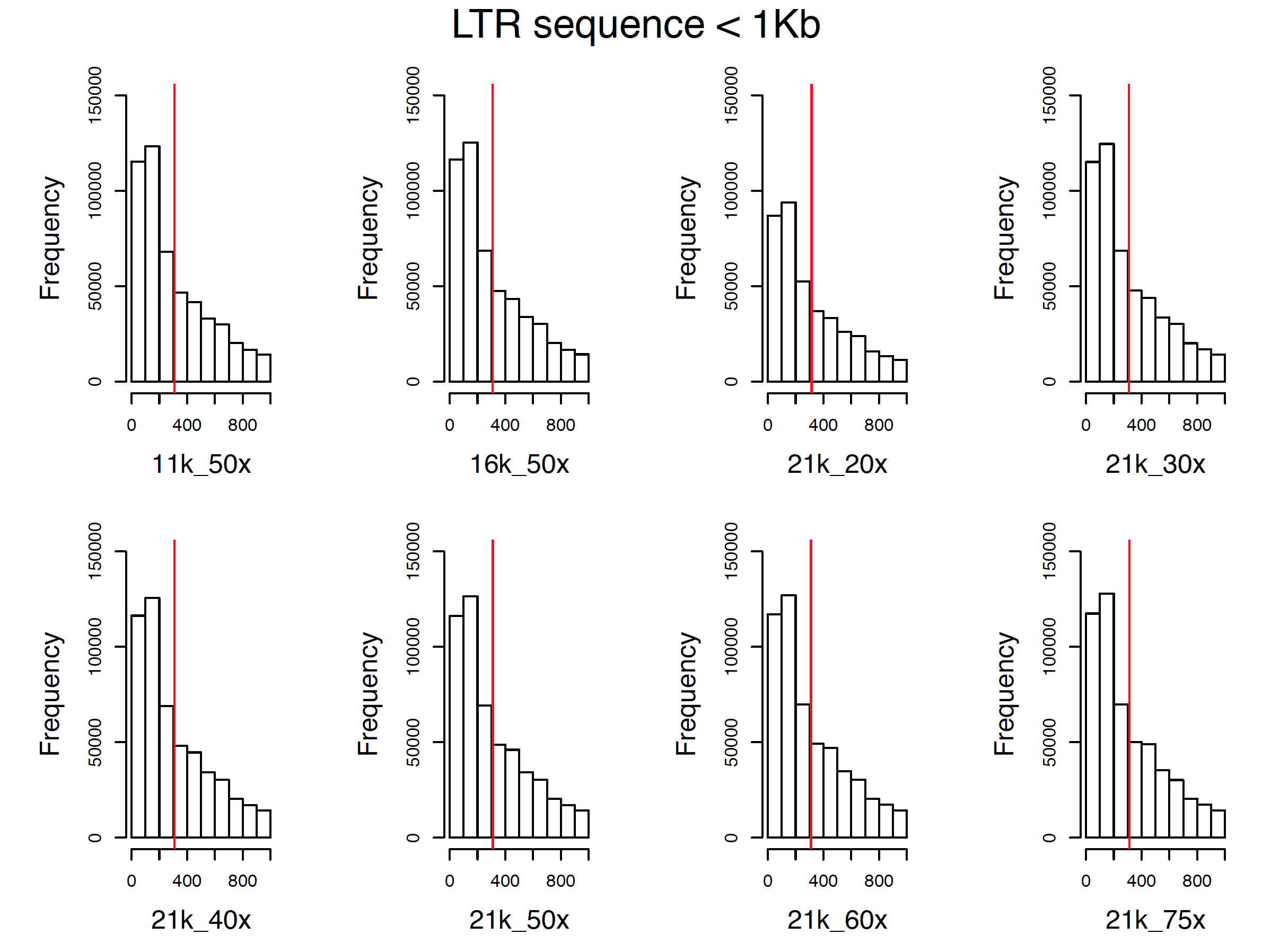

Figure S10. Length distribution of fragmented LTR retrotransposons shorter than 1 kb in NC358 assemblies. X-axes, length of LTR fragments. Y-axes, count of LTR retrotransposons. Red lines, mean length of LTR retrotransposons.

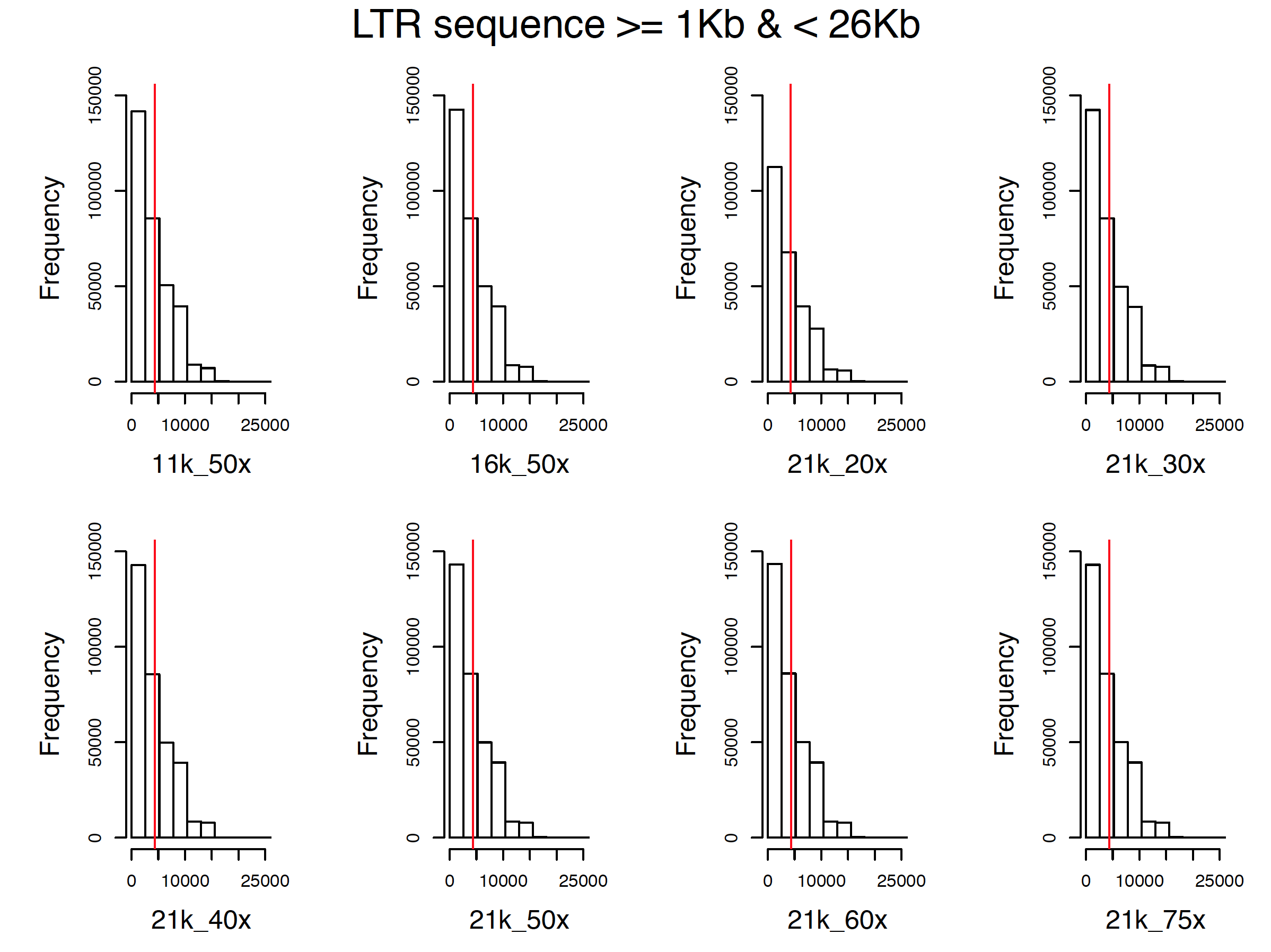

Figure S11. Length distribution of fragmented LTR retrotransposons ranging from 1 kb to 26 kb in NC358 assemblies. X-axes, length of LTR fragments. Y-axes, count of LTR retrotransposons. Red lines, mean length of LTR retrotransposons.

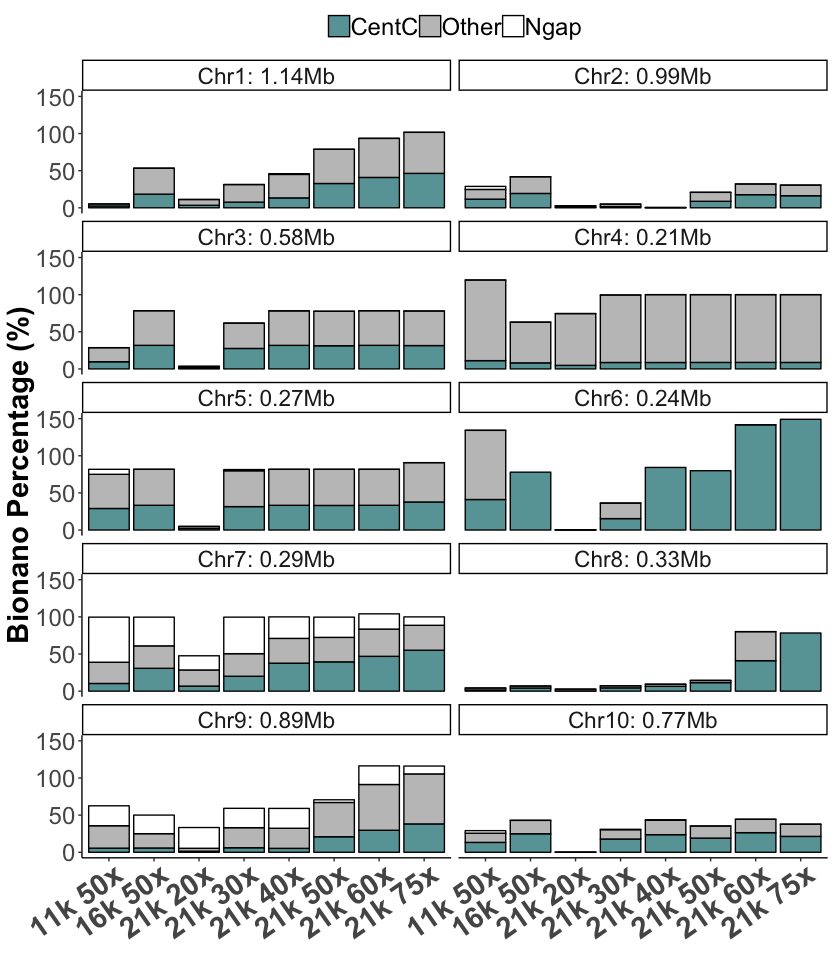

Figure S12. Evaluation of CentC array assembly using the bionano optical map. CentC, CentC array found in the ChIP-defined centromere region. Ngap, non-13N gaps. Other, other sequence contents.

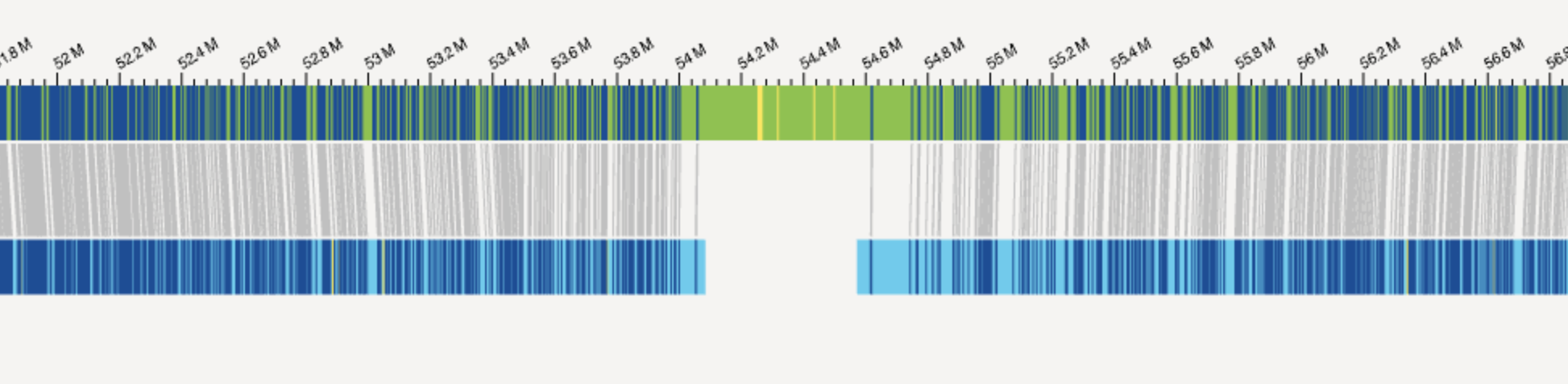

Figure S13. An example of failure in Bionano *de novo* assembly due to lack of DLE-1 label in CentC regions, where sequence assembly could be more contiguous. The upper panel represents the reference genome, which is the pseudomolecule chr6 in the 21k_75x assembly. The lower panels are the Bionano *de novo* assembled optical maps used as queries. The blue bars indicate the matching label sites between reference and query, while yellow bars represent unmatched sites.

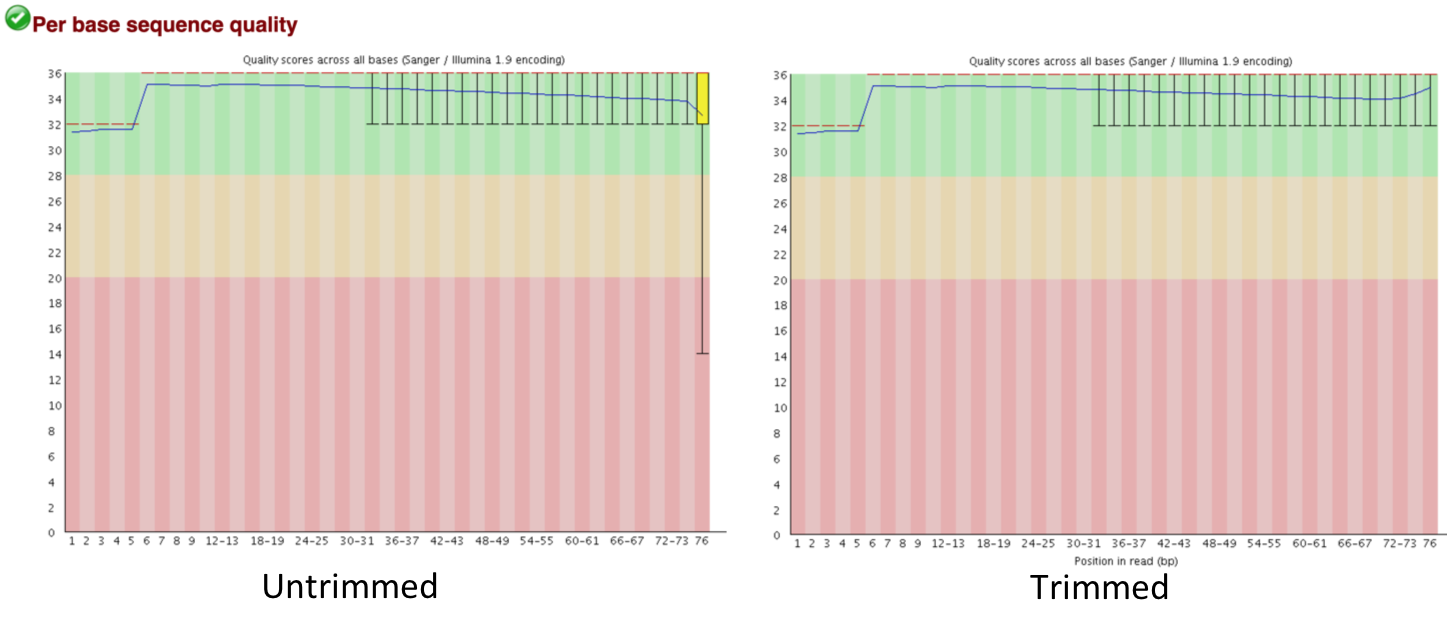

Figure S14. An example of RNA-seq libraries for quality trimming. X-axes, base pairs on RNA-seq reads. Y-axes, Phred-scaled quality scores.

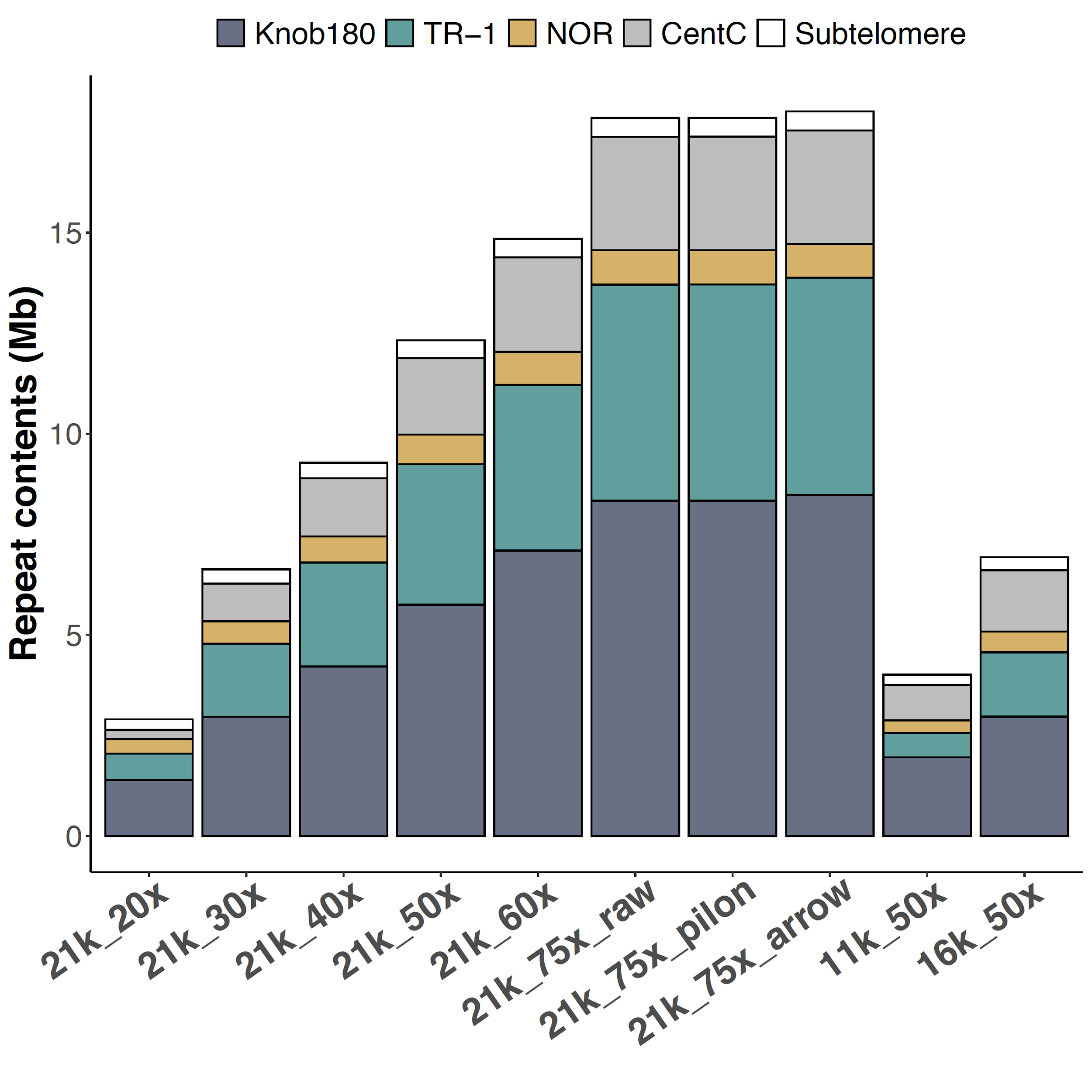

Figure S15. Assembled repeat contents in NC358 assemblies. The 21k_75x_pilon assembly was polished using Pilon with 30x Illumina short reads. The 21k_75x_arrow assembly was polished using Arrow with the full 75x PacBio dataset. The rest of NC358 assemblies are non-polished.

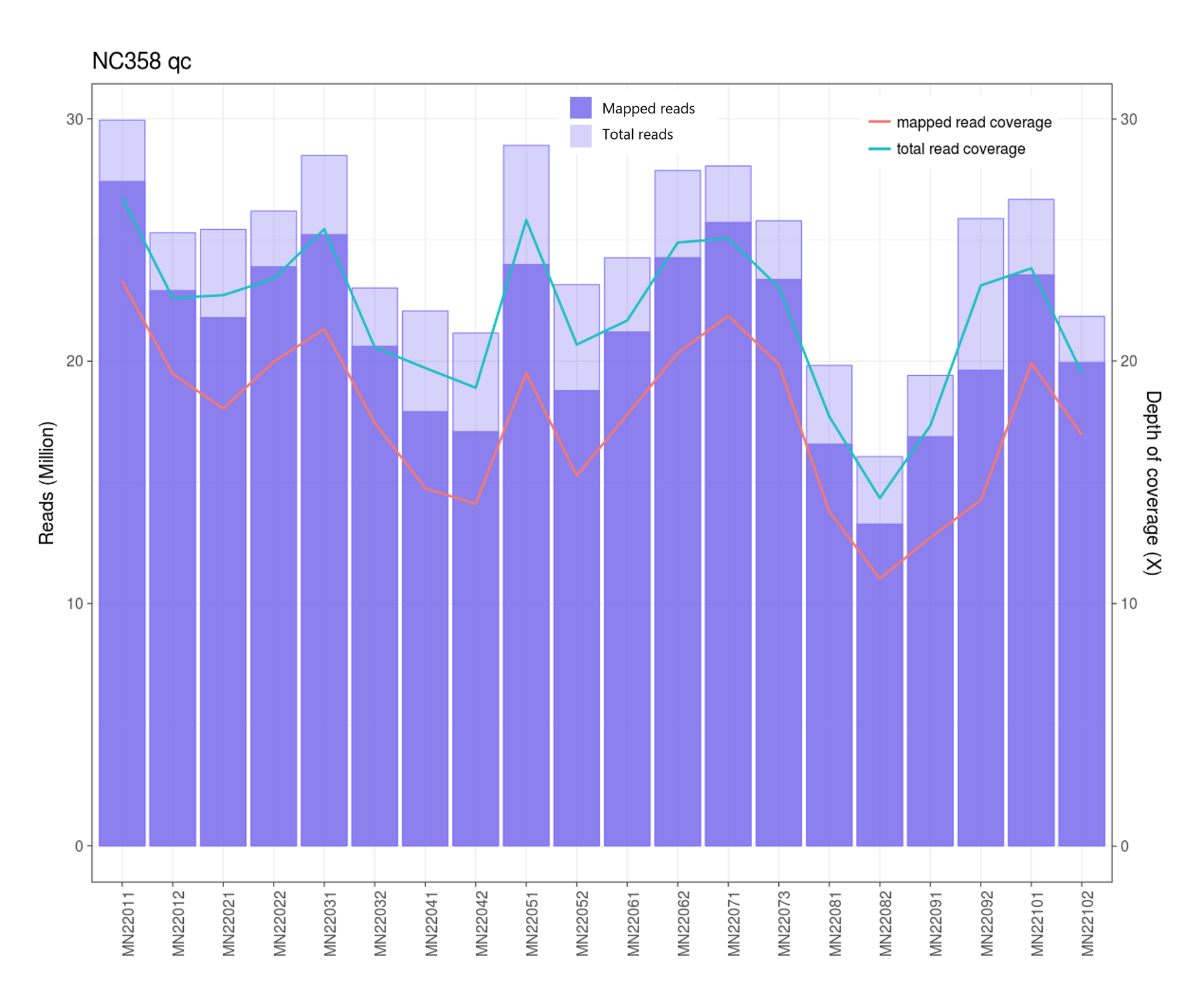

Figure S16. Quality checks of NC358 RNA-seq libraries. RNA sequences were trimmed and mapped to the B73 v4 genome. Read coverage was calculated using a total CDS length of

266 MB. MN22031, V11 base; MN22032, V11 base; MN22021, 8DAS shoot; MN22022, 8DAS shoot; MN22062, V18 tassel; MN22061, V18 tassel; MN22051, V11 tip; MN22052, V11 tip; MN22041, V11 middle; MN22042, V11 middle; MN22081, R1 anther; MN22082, R1 anther; MN22102, 16DAP embryo; MN22101, 16DAP embryo; MN22073, V18 ear; MN22071, V18 ear; MN22012, 8DAS root; MN22011, 8DAS root; MN22092, 16DAP endosperm; and MN22091, 16DAP endosperm.
